# Accurate and automated high-coverage identification of chemically cross-linked peptides with MaxLynx

**DOI:** 10.1101/2021.08.26.457759

**Authors:** Şule Yılmaz, Florian Busch, Nagarjuna Nagaraj, Jürgen Cox

## Abstract

Cross-linking combined with mass spectrometry (XL-MS) provides a wealth of information about the 3D structure of proteins and their interactions. We introduce MaxLynx, a novel computational proteomics workflow for XL-MS integrated into the MaxQuant environment. It is applicable to non-cleavable and MS-cleavable cross linkers. For both we have generalized the Andromeda peptide database search engine to efficiently identify cross-linked peptides. For non-cleavable peptides, we implemented a novel di-peptide Andromeda score, which is the basis for a computationally efficient N-squared search engine. Additionally, partial scores summarize the evidence for the two constituents of the di-peptide individually. A posterior error probability based on total and partial scores is used to control false discovery rates. For MS-cleavable cross linkers a scoring of signature peaks is combined with the conventional Andromeda score on the cleavage products. The MaxQuant 3D-peak detection was improved to ensure more accurate determination of the monoisotopic peak of isotope patterns for heavy molecules, which cross-linked peptides typically are. A wide selection of filtering parameters can replace manual filtering of identifications, which is often necessary when using other pipelines. On benchmark datasets of synthetic peptides, MaxLynx outperforms all other tested software on data for both types of cross linkers as well as on a proteome-wide dataset of cross-linked *D. melanogaster* cell lysate. The workflow also supports ion-mobility enhanced MS data. MaxLynx runs on Windows and Linux, contains an interactive viewer for displaying annotated cross-linked spectra and is freely available at https://www.maxquant.org/.

## Introduction

Chemical cross linking combined with mass spectrometry has undergone remarkable developments to become a promising complementary method for studying protein structure, conformation and interactions^1–4^. A typical protein cross-linking experiment starts with forming covalent bonds between spatially close residues in proteins or protein complexes through a cross linker. A chemical cross linker is typically specific to certain amino acids and its presence imposes a distance constraint^5^. A cross linked protein sample is then enzymatically digested and the resulting complex peptide mixture contains different types of products, including mono-, loop- and cross-linked peptides^6^. In fact, cross linked peptides can be more diverse than just two peptides connected by a single linker due to combination of multiple reactions, resulting in higher-order cross linked peptides^7 8^. Linear peptides, in which cross linkers are not attached, are usually much more abundant in the sample than cross-linked products and therefore enrichment of cross-linked peptides is generally required. The peptide mixture is analyzed by liquid chromatography coupled to tandem mass spectrometry (LC-MS/MS) and the resulting experimental MS/MS spectra are assigned to cross linked peptides by using specialized algorithms^4,9^. Eventually, the cross linked peptide identifications are used to gain insights into protein structures or their interaction partners^10,11^.

XL-MS peptide identification algorithms can be sub-divided based on the type of the supported cross linker, which can be non-cleavable or MS-cleavable. A non-cleavable cross linker remains intact during mass-spectrometric analysis, whereas an MS-cleavable cross linker fragments easily owing to a labile bond. MS/MS spectra produced in these two approaches are qualitatively different. In both cases, the identification algorithm has to include different crosslink products in addition to single peptides to the search space, strongly increasing its complexity compared to conventional proteomics. In studies with non-cleavable cross linkers, any peptide can be linked to any other peptide which causes a search space increase, known as the n-squared problem^1,6^. Many programs like StavroX^12^ and OpenPepXL^13^ use an exhaustive search in that space, which is computationally challenging and therefore non-cleavable cross linkers are commonly used for smaller sets of proteins or protein complexes. Some other algorithms have different approaches to tackle this search space problem. For example, xQuest^14^ detects isotopic pairs on the MS1 level with a mass shift observed by using of the heavy- and light-labeled cross-linked peptides to select candidate peptides, and by that reduced the number of MS/MS spectra that need to be submitted to the search. Kojak^15^ has a two-pass approach: the first pass selects peptide candidates while allowing a differential modification mass and the second pass creates cross-linked peptides from these candidates. In addition to this search space challenge, non-cleavable cross-linked peptides can show an unequal distribution of the fragment ions from the two peptides^6^. These two issues can be circumvented by using MS-cleavable cross linkers^16^, which enable even proteome-wide applications of XL-MS^17,18^. The fragmentation of the labile bond of the MS-cleavable cross linker produces single peptides with specific parts of the cross linker attached. This results in the observation of distinctive signature ions^16^. With the help of the signature peaks and the corresponding precursor, candidate cross-linked peptides are produced and then scored against the experimental spectrum. XlinkX^17^ and MeroX^19^ are two commonly used algorithms in MS-cleavable studies that use signature peaks in their algorithms.

MaxQuant^20^ is a freely available computational proteomics software widely used in the community that supports diverse experimental designs and mass spectrometry platforms. Here we describe the integration of novel tools and algorithms for the identification of cross-linked peptides into MaxQuant, collectively called MaxLynx. We evaluated MaxLynx on synthetic peptide data sets of cross-linked peptides obtained with the non-cleavable and MS-cleavable approaches and compared its performance to results obtained with several other software packages. We found that at 1% false discovery rate (FDR), MaxLynx outperformed many other software for both non-cleavable and MS-cleavable data sets with up to four times more cross-linked-to spectrum-matches (CSMs) and twice the number of unique cross links. In addition, we performed a complex proteome-wide study and compared that to the published results from the existing algorithm. We observed that MaxLynx again reported more CSMs along with more unique cross-links.

## Experimental section

### Synthetic benchmark dataset

We downloaded the raw data from the dataset with identifier PXD014337^21^ in the PRIDE repository^22^, which contains cross linked synthetic peptides linked by non-cleavable and MS-cleavable cross linkers and measured via LC-MS/MS. In total, 95 tryptic peptides from the *S. pyogenes* Cas9 protein were chemically synthesized. Each of these peptides contains one internal lysine residue for cross-linking. Both the peptide N- and C-termini were modified to prevent unwanted crosslinking reactions. These peptides were split into 12 groups and cross-linking experiments were performed only within each group. These 12 samples were then mixed before introduction to LC-MS/MS (Orbitrap Q-Exactive HF-X). Disuccinimidyl suberate (DSS) was used to create non-cleavable cross-linked peptides and these were measured as three technical replicates. Disuccinimidyl dibutyric urea (DSBU) and disuccinimidyl sulfoxide (DSSO) were used to create MS-cleavable data and measurements were done with stepped higher-energy collision-induced dissociation (HCD) on an Orbitrap Q-Exactive HF-X instrument without technical replication for each data set.

### Proteome-wide benchmark dataset

We downloaded the raw data from dataset PXD012546^18^ in the PRIDE repository to evaluate the performance of MaxLynx in proteome-wide studies. Three biological replicates of *D. melanogaster* (fruit fly) embryo extracts were cross linked by using DSBU and separated with size exclusion chromatography resulting in 67 LC-MS/MS runs. The samples were measured with stepped HCD on an Orbitrap Q-Exactive Plus mass spectrometer^18^.

### timsTOF Pro BSA dataset

Bovine serum albumin (GERBU Biotechnik GmbH, #1062) was dissolved in 50 mM PBS pH 7.0 and the protein solution was transferred into Amicon Ultra-0.5 mL centrifugal filters (10 kDa NMWCO). After several rounds of dilution with 50 mM PBS pH 7.0 and re-concentration by centrifugation to remove potential interfering small molecules, the protein concentration was adjusted to 10 µM. Proteins were cross linked at a molar ratio of cross linker to protein of 25 : 1 by addition of 1 µL 50 mM DSSO (ThermoFisher Scientific, #A33545) and 50 mM DSBU (Bruker Daltonics, #1881355) in DSMO, respectively. After overnight reaction at 4 °C, the reactions were quenched by addition of 100 µL 100 mM Tris/HCl pH 7.5. Proteins were denatured by buffer-exchange to 50 mM ammonium bicarbonate with 8M urea, and reduced/alkylated by incubation with DTT (5 mM) for 30 min and iodoacetamide (15 mM) for 20 min at room temperature. Proteins were buffer-exchanged to 50 mM ABC and digested with 4 µg Trypsin Gold (Mass Spectrometry Grade, Promega, #V5280) for 1 h at 37 °C. The flowthrough after centrifugation was combined with the flowthrough from an additional wash of the filters with 200 µL water with 0.1% formic acid. After evaporation of solvent by SpeedVac Vacuum concentration, peptides were re-suspended in water with 0.1% formic acid. *LC-MS/MS*. 200 ng digested sample was analyzed with a NanoElute coupled to a timsTOF Pro mass spectrometer (Bruker Daltonics). Peptides were separated on an Aurora C18 column (25 cm x 75 µm ID, 1.6 µm particle size, Ionoptics, AUR2-25075C18A-CSI) at a flow rate of 0.4 µL/min at 50 °C. The following gradient was used: within 60 min from 2 to 17% B, within 30 min from 17 to 25% B, within 10 min from 25 to 37% B, within 10 min from 37 to 80% B, isocratic at 80% B for 10 min. Solvent A was water with 0.1% formic acid and solvent B was acetonitrile with 0.1% formic acid. Mass spectra were recorded from m/z 100 -1700 and inverse reduced mobility of 0.6 - 1.52 Vs/cm2. Charge states for PASEF were set to 3-5 and selected ions were fragmented by TIMS stepping with two 1/K0-dependent collision energies (collision energies linearly interpolated between 0.85 – 1.2 Vs/cm2 to 25 – 55 and to 30 – 70 eV). 10 PASEF MS/MS scans were triggered per cycle (2.23 sec).

### Data analysis

#### Synthetic benchmark dataset

For the synthetic cross-linked peptide library datasets (PXD014337), the common search settings were appended to the given table given by Beveridge and co-workers^21^ (See Table S1 and Table S2). The search databases used were Cas9 plus 10 proteins and Cas9 plus 116 cRAP contaminants proteins for the non-cleavable and MS-cleavable data sets, respectively. These default MaxQuant search settings were changed: ‘Include contaminants’ was disabled and including any loss and also higher charge states in the MS/MS analyzer section were also disabled. The new option called ‘Peak Refinement’ was enabled. No cross link-specific filtering was enabled, which are the minimum score for cross-linked peptides, the minimum score for other cross linked products and the minimum number of fragment ions from each peptide. The minimum partial score remained as the default value of 10 for both the non-cleavable and the MS-cleavable data sets. For all data sets, the MaxQuant results were kept at the cross-linked peptide-to-spectrum matches false discovery rate (CSM-FDR) of 1% the further comparison. Because of the peptide level experimental design^21^, the software performance was evaluated without relying on protein structures: from a given list of CSMs, a CSM was considered to be correct when cross linked peptides belonged to the same peptide group, otherwise it was assigned as incorrect.

#### Proteome-wide benchmark dataset

The following settings were changed from MaxQuant default values: a minimum peptide length of 5 and max peptide mass 8000 Da was used. No contaminants were added. Only cross-linked peptides (inter-peptides with a single cross link modification) were considered to be able to fairly compare against the published results. The selected enzyme was trypsin with four missed cleavages. The peptide and fragment ion tolerances were 5 ppm and 15 ppm. No high-charges or losses were allowed for the FTMS MS/MS analyzer settings. Carbamidomethylation of a cysteine was chosen as a fixed modification and oxidation of a methionine and acetylation of protein N-terminus were as a variable modification with allowing two modifications per peptide. The search database contained the identified *Drosophila* proteins (ID with 9535 protein entries), as was recommended by Götze and co-workers^18^. The MaxLynx results were compared at 1% FDR.

#### tims-TOF Pro BSA dataset

The search database contains the bovine serum albumin protein plus *Pyrococcus furiosus* (PFU) proteins (502 reviewed protein sequences, downloaded on March 8^th^, 2021 from UniProtKB^23^). Using PFU proteins as an entrapment database was previously proved as a good way to validate proteomics results^24^. In our analysis, we used this approach to evaluate the performance of MaxLynx. A CSM including only BSA proteins was likely to be correct whereas any CSM to PFU proteins was incorrect. This additional criterion further allowed us to validate our cross-link identifications independent of protein structure information. The search settings were as follows. Oxidation of methionine with and carbamidomethylation of cysteine is taken as variable and fixed modification respectively with a maximum of one variable modification per peptides. Trypsin is selected as enzyme with maximum three missed cleavages. The cross linkers are either DSSO or DSBU (both as heterobifunctional as a lysine residue can be linked to lysine, serine, threonine and tyrosine residues) with mono- and cross-linked peptides options. The minimum peptide length was six and the maximum peptide mass 6000 Da. Contaminants were not included. The default MaxQuant maximum charge was changed from 4 to 6 in the Bruker TIMS instrument settings. Higher charges and losses addition were disabled for the MS/MS analyzer.

### Software availability, requirements and usage

MaxLynx is freely available at www.maxquant.org as part of MaxQuant, The crosslink search module was written in the C# programming language, and integrated into the existing MaxQuant workflow^20^. The program can run as a graphical user interface (GUI) tool on Windows and also on the command line (CLI) on both Windows and Linux operating systems.

## Results and discussion

### The MaxLynx workflow

Much of the MaxQuant workflow for conventional peptides is used in MaxLynx as well (Figure 1a), as for instance, the detection and de-isotoping of features in the MS1 data. These are usually three-dimensional objects spanned by m/z, retention time and signal intensity. We also support ion-mobility enhanced data^25^, generated from timsTOF Pro instrument, in which case the MS1 features become four-dimensional. The novel peak refinement feature (Figure 1b) is executed after the peak detection for data without ion mobility and assembles peaks that were not properly put together due to noise, which may happen for peptides with higher mass. These not well-assembled peaks lead to wrong assignments of the monoisotopic peak to the isotope pattern^26^ which in turn hinders their identification. This problem is strongly reduced by the peak refinement. The conventional Andromeda search engine^27^ is used to identify linear peptides and also to perform nonlinear mass recalibration. For the identification of cross-linked peptides, one of two specialized search engines is used (Figure 1c), depending on the cross linker being MS-cleavable. Finally, a module for applying a desired false discovery rate on the level of CSMs based on a posterior error probability calculation is included. A list of all detailed steps involved in MaxLynx is shown in Figure S1. A user’s guide on how to use MaxLynx can be found in the Supporting Information. Any bi-functional non-cleavable or MS-cleavable cross linker is configurable in the user interface (Figure S2).

**Figure 1.**
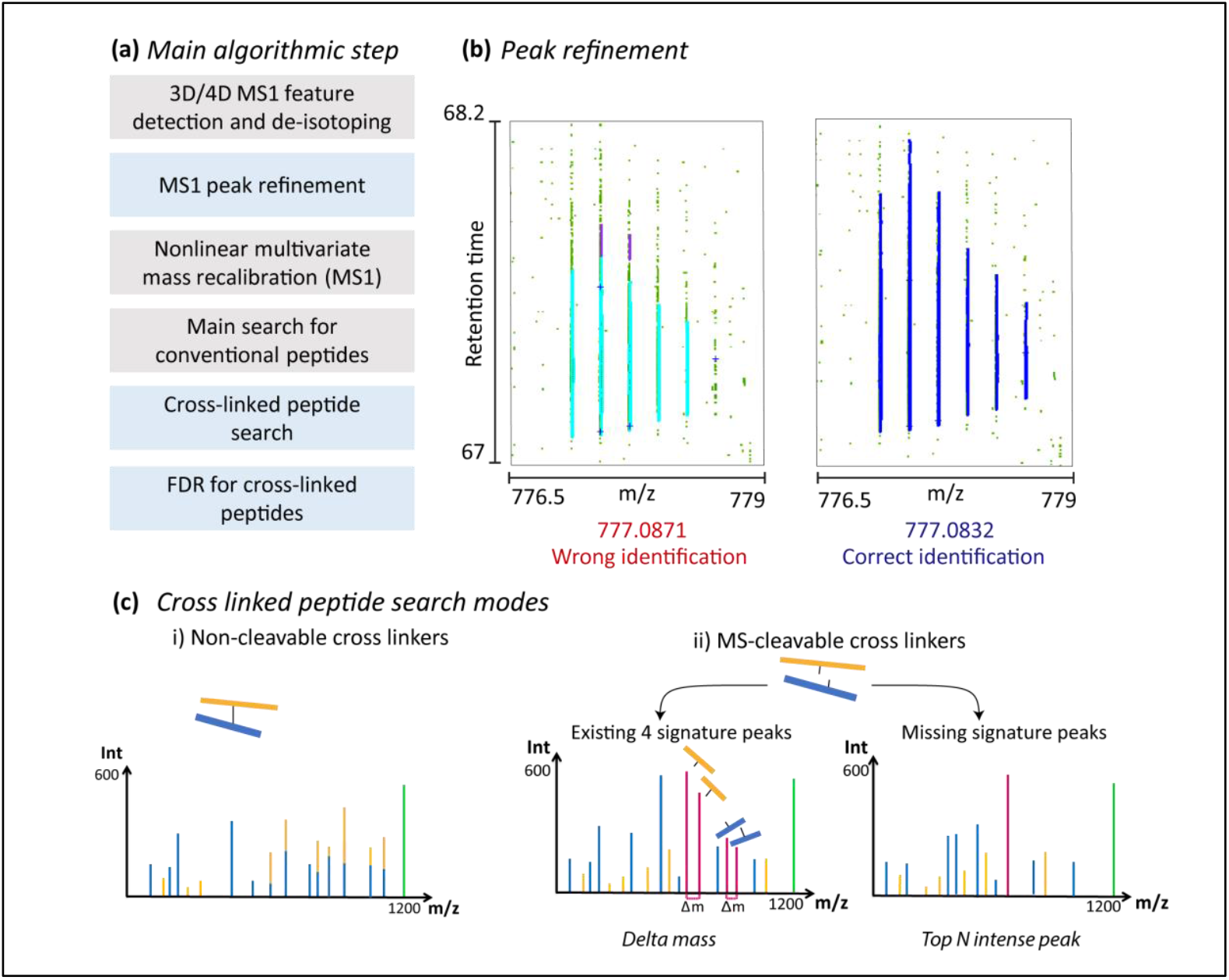
The computational workflow of MaxLynx. (a) Schematic simplified block diagram of the main algorithmic steps involved in MaxLynx. Steps in grey are unchanged from the MaxQuant workflow for regular peptides while blue steps are newly developed for the cross-linking search. (b) Peak refinement is a computational step which was added after the peak detection with the aim of ‘repairing’ peaks typically of heavy mass that are not well defined due to noise. (c) Depending on the linker that has been applied is MS-cleavable, one of two search engines is employed to query the measured MS/MS spectra.

Cross-linked peptide identification results can be further inspected through the MaxQuant Viewer interactively when a MaxLynx run is successfully completed (Figure 2). All defined cross link products such as mono-link or di-peptides can be viewed, also for ion-mobility enhanced data sets. For di-peptides, the shorter (beta) and longer (alpha) peptides are color coded same on both ‘Spectrum’ and ‘Peptide Sequence’ panels. On the ‘Spectrum panel’, ion types, its number and its m/z values are provided. On the ‘Peptide Sequence’ panel, a cross linker is shown in red between two peptides (See Supporting Information).

**Figure 2.**
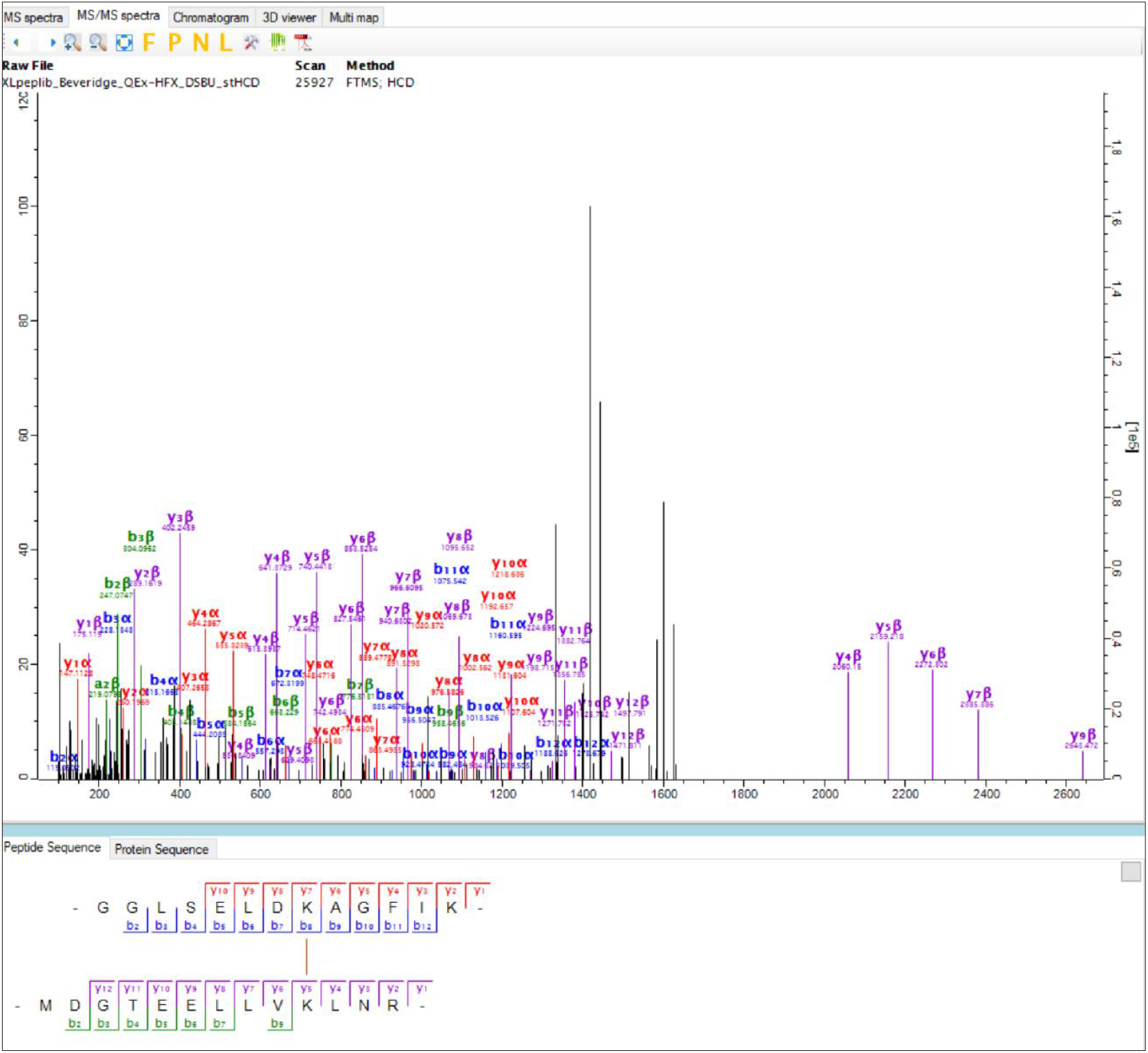
Visualization of identified cross-linked peptide. A screenshot of the MS/MS viewer in MaxQuant with annotation for fragments resulting from cross-linked peptides.

### Non-cleavable cross-linked peptide search

MaxLynx generates a complete search space for non-cleavable cross-linked peptides and performs an exhaustive search in it. The first step is to generate *in silico* peptides depending on the Andromeda search settings and then the search space is constructed by combining all putative peptides. The cross-link search space includes single peptides and different cross-linking products (di-peptides, mono-linked or loop-linked peptides) in case a peptide has a target residue for the cross linker of choice. A user can define cross-linking products on the new group specific parameter panel called ‘Cross links’ (See Supporting Information and Figure S3). Mono-linked peptides are added to the MaxLynx searches because two adjacent peptides can mimic a mono-linked peptide or di-peptides since they have the same precursor mass^28^. In case there are two or more target resides on a peptide, loop-linked peptides are created. In addition to these three cross-linked peptide products, single peptides without cross-linker modifications are added to the search space as well by default. Furthermore, MaxLynx can consider higher-order cross linked peptides, which are the combination of different single crosslink modifications (e.g. an inter-peptide product has also a mono-link, also known as Type2,1^7^). Note that the complexity of the search space will be increased even more when these cross-linking products are added, and hence the results will include more false positives. Therefore, many other algorithms do not consider such multiple modifications. The cross linking products, which are mono-, loop- and cross-linked, and if selected higher-order cross linked peptides, are internally written to temporary files with their masses, the number of links and the linked sites and peptide level information. Then for each crosslink product, variable modifications are added to their masses and then a single index is created^27^.

The second main step after the cross-link space construction is the MS/MS crosslink search. This step is integrated into the established Andromeda scoring workflow. The precursor mass of an experimental MS/MS spectrum is compared against indexed masses and when an indexed mass is equal to the experimental precursor mass within a certain tolerance, a theoretical spectrum is generated. It is important to highlight the main difference compared to the ordinary peptide searches: a theoretical spectrum for cross-linked peptides has theoretical fragment ions from both peptides and ions from the residue contributing to cross-linking have a corresponding mass shift.

### MS-cleavable cross-linked peptide search

Another approach to study cross-linked peptides is based on using MS-cleavable cross linkers. During MS/MS analysis, the cross linker is cleaved and this cleavage results in two peptides with partial cross linkers attached (Figure 1c). The longer of the two peptides is denoted with the Greek letter alpha, the shorter with beta. The structures of the various MS-cleavable cross linker molecules vary but typically they have two labile bonds so the cleavage commonly results in peptides containing either a shorter or a longer piece of the cross linker and this produces in the MS/MS spectrum a characteristic doublet peak-signal per peptide with a specific mass difference. With the help of the doublet signals, the masses of the two peptides can be determined individually and cross-linked peptides are identified based on this knowledge. In MaxLynx, we consecutively apply three approaches to detect signature peaks, the strict mass difference approach, the top intensities approach and finally a second round of the mass difference approach with relaxed criteria. The strict mass difference approach depends on observing mass differences between the long and short versions of the remainder of the cleaved cross linker on the same peptide for both pairs of peaks in the MS/MS spectrum. The top intensities approach checks for the most intense peaks in the MS/MS spectrum if it can be interpreted as one of the signature peaks, without requiring the other signature peaks to be present. In the mass difference approach with relaxed criteria only one pair of signature peaks is required. All three approaches work on the MaxQuant processed version of MS/MS spectra for which peaks have been de-isotoped and higher charge states of fragment ions have been transformed to charge one^27^.

In the strict mass difference approach, we aim to find all four signature peaks with two pairs having the expected mass difference between the long and short linker residual denoted as Δm.. For that purpose, we loop through all peaks in the MS/MS spectrum that are larger than a user definable minimal mass with the hypothesis that it is the β-peptide with the shorter linker residual (βs). Then we check for the presence of the three corresponding remaining signature peaks, which are the β-peptide with the longer linker residual (βl) and both versions of the longer peptide (αs and αl) whose masses are given by

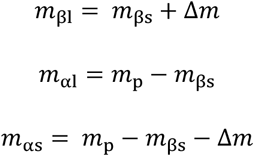

where *m*_p_ is the measured precursor mass of the whole di-peptide. In case all four peaks are found with the given precursor mass tolerance they are accepted as the signature peaks. Based on this, all theoretical spectra are constructed with y- and b-ion series for linear α and β peptides whose masses are compatible within the mass tolerance.

A weakness of the mass difference approach is that four signature peaks must be observed. However, it is not always the case that all of these are present in the spectrum^29^. Furthermore, there could be homo-dimeric peptides, meaning that the peptide is linked to a peptide with the same sequence and therefore only two signature peaks exist. Indeed, a recent study^21^ showed that XlinkX^17^, an algorithm using the mass difference strategy, did not report any such homo-dimeric peptides. To overcome this problem, we implemented a second step to find signature peaks in a less stringent way by choosing candidate peaks based on their intensities^29^. This is done whenever the mass difference approach does not find a solution. The assumption here is that the signature peaks are among the most intense peaks^29^. We go through the top *n* most intense peaks in the MS/MS spectrum, where *n* = 3 by default which can be changed by the user. For each of the top peaks it is hypothesized that it is either carrying the longer or the shorter linker residue. Knowing the precursor mass, both hypotheses lead to masses for the α and the β peptide. If both the calculated α- and β-peptide masses are heavier than the given minimum peptide mass, a theoretical combined spectrum is submitted to the database search. We observed that the signature peaks could sometimes have an ammonia (NH_3_) loss. Therefore, we extended the assumption above by considering that the top intense peaks can also have such a loss. This is expected especially for fragments containing lysine residues^27^. Based on the α and β peptides found, cross linked peptide products are *in silico* constructed followed by theoretical MS/MS spectrum generation and scoring in the Andromeda search^27^. In case neither of the two approaches described finds a candidate explanation for an MS/MS spectrum, we perform another round with the mass difference approach in which it is sufficient if only one peak pair with the characteristic mass difference is found.

### False discovery rate control

The FDR control in MaxLynx is based on the target-decoy strategy. A cross link search results in a list of cross-linked-peptide-to-spectrum matches (CSMs) which can be split into three crosslink product groups: i) di-peptides, in which two peptides are linked ii) single peptides with attached linker molecule, which are mono- and loop-linked peptides and iii) single ordinary peptides that no cross linker is attached to. Di-peptides can be target-target (TT), target-decoy (TD) or decoy-decoy (DD) cross-linked di-peptides.

The posterior error probability (PEP) is the likelihood of a CSM being wrongly identified at a given Andromeda score and additional selected peptide properties. The PEP calculation by MaxQuant^20^ includes the logarithm of an identified peptide length. For MaxLynx, however, we modified our implementation to calculate PEP for di-peptides, as a consequence of the co-existing two peptides. An unequal fragmentation is problematic for di-peptides^30^ and has a negative effect on scoring algorithms because search engines can still assign a good score despite very little or no evidence from one peptide but good fragmentation on its paired peptide. To overcome this issue, we decided to use the minimal partial score instead of peptide length at our PEP calculation for di-peptides. The partial score is a version of the Andromeda score that is calculated using only ions from one peptide of the cross-linked di-peptide against a given experimental MS/MS spectrum. Every cross-linked di-peptide has two partial scores, an α and a β partial score, and out of these the minimum is used for the PEP calculation. The PEP is subsequently corrected for the number of modifications, the precursor charge state and the biggest number of missed cleavages from the peptides involved in di-peptides. Afterwards, all CSMs are sorted based on their PEP scores in an ascending order and the FDR is calculated as a number of false CSMs divided by a number of target CSMs.

### Benchmarks on synthetic cross-linked peptides

We re-analyzed publicly available data sets, in which synthetic peptides were cross-linked with the non-cleavable cross linker DSS and with the MS-cleavable cross linkers DSSO and DSBU. Beveridge and co-workers^21^ benchmarked these data sets by using several existing software platforms. For the non-cleavable cross linker data pLink2^31^, StavroX^12^, Xi^32^ and Kojak^15^ were benchmarked, whereas for the MS-cleavable cross linker data MeroX^19^ and XlinkX^17^ were used. This data was also analyzed with OpenPepXL in their own publication^13^. We compared the MaxLynx results to the results provided in these two studies in terms of the number of CSMs and unique cross-linking sites at CSM FDR=1%.

For the non-cleavable cross linker data set, MaxLynx reported the highest number of CSMs at FDR=1% compared to the other algorithms with 839 correct and 12 incorrect CSMs on average (Figure 3 and Tables S3-S4). Moreover, MaxLynx reported the highest number of unique cross-links (on average 216). In non-cleavable cross linker search, the settings that influence the number of the identification the most was related to the peak refinement option. When this option was disabled, the average number of correct CSMs dropped to 666 and the average number of correct cross-links to 199. This shows that the studies on the improvement of feature detection for heavier peptides has a significant improvement on cross-linked peptide identifications. We recommend the peak refinement option to be on for cross-link searches.

**Figure 3.**
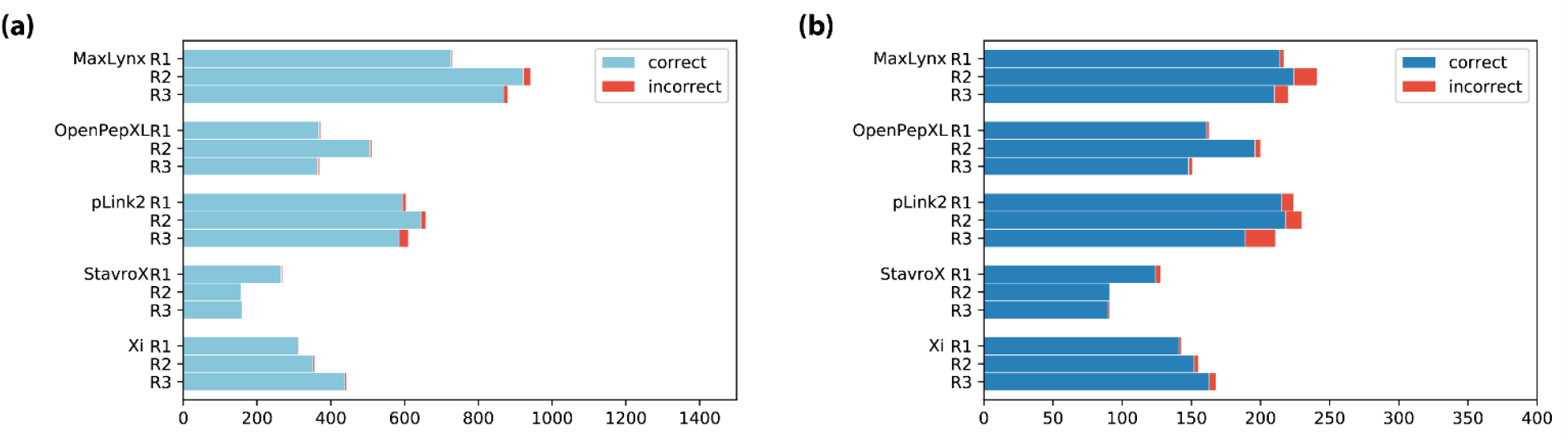
Comparison of MaxLynx against other cross link search engines on the non-cleavable cross-linker dataset. (a) shows the number of cross-linked-to-spectrum matches (b) shows the number of unique cross links at FDR=1%. The results of OpenPepXL were obtained from Netz and co-workers^13^ and the results of the other software were taken from Beveridge and co-workers ^21^. Three replicates were shown as R1, R2 and R3.

On the MS-cleavable cross-linker data set, MaxLynx reported the highest number of correct unique cross links for both DSBU and DSSO data sets compared to the most of the other search engines^21^ (Figure 4 and Tables S5). For the DSBU data set, MaxLynx resulted in 240 correct and 12 incorrect unique crosslinks at FDR=1%. The MaxLynx results on the DSSO data set showed a similar pattern as the DSBU results. MaxLynx reported the highest number of correct crosslinks at FDR=1%, with 185 correct and 10 incorrect unique crosslinks. One reason why MeroX performed better in terms of identified number of cross-links was the ability to detect homo-multimeric inter-peptides^21^, in which the peptide is linked to a peptide with the same sequence. For MaxLynx, the number of correct cross-links coming from the CSMs of homo-multimeric inter-peptides was 48 and 41 for the DSBU and DSSO, respectively. When these homo-multimeric cross-links are ignored (192 and 144), MaxLynx still identifies a higher number of cross-links than are found with XlinkX.

**Figure 4.**
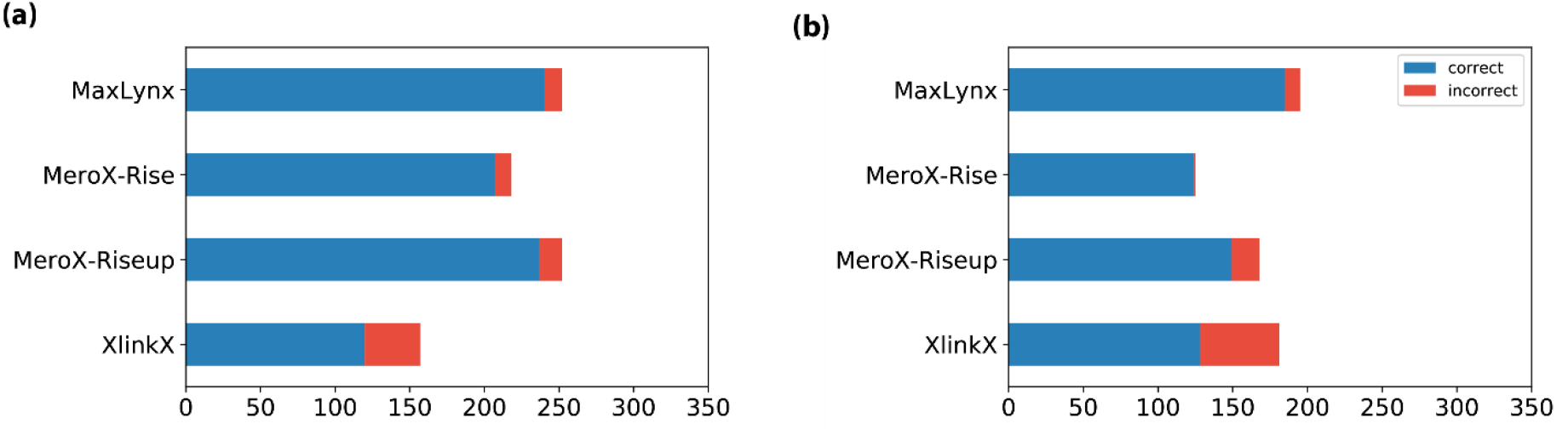
Comparison of MaxLynx against other cross link search engines on the MS2 cleavable cross-linker datasets. (a) and (b) show the number of unique cross links respectively at the DSBU and DSSO data sets at FDR=1%. The results of the other search engines were obtained from Beveridge co-workers^21^.

### Benchmark on proteome-wide MS-cleavable cross linker data

Next, we evaluated the capability of MaxLynx to analyze large-scale, proteome-wide cross linking data sets. For that purpose, we re-analyzed the PRIDE dataset PXD012546 of *D. melanogaster* embryo extracts cross linked with DBSU and compared against the published results. At FDR=1%, MaxLynx reported in total 34,819 CSMs and 7,579 unique cross links exceeding the originally reported number by MeroX while using the same settings (Table 1). While the reproducibility of identification results between the three replicates is with 18% (Figure 5a) found in all three rather low, it is higher than reported with Merox (overlap of 15%). As noted by Götze and co-workers^18^ the reasons for this observation could be attributed to experimental and biological conditions. Next, we checked the number of unique cross links overlapping between MaxLynx and MeroX software and we observed that around one-third of all found unique cross links were shared between these two (Figure 5b).

**Table 1.** The overview of the results for the proteome-wide study at FDR=1%. MeroX results were obtained from Götze and co-workers^18^. Note that the number of unique cross links and unique intra-molecule cross links were directly taken from their supplementary data and their public PXD012546 data set.

| Software | #CSMs | #unique cross-links | #unique intra-molecule cross-links | #unique inter-molecule cross-links |
| --- | --- | --- | --- | --- |
| MeroX | 29,931 | 7,428 | 5,817 | 1,611 |
| MaxLynx | 34,819 | 7,579 | 6,234 | 1,345 |

**Figure 5.**
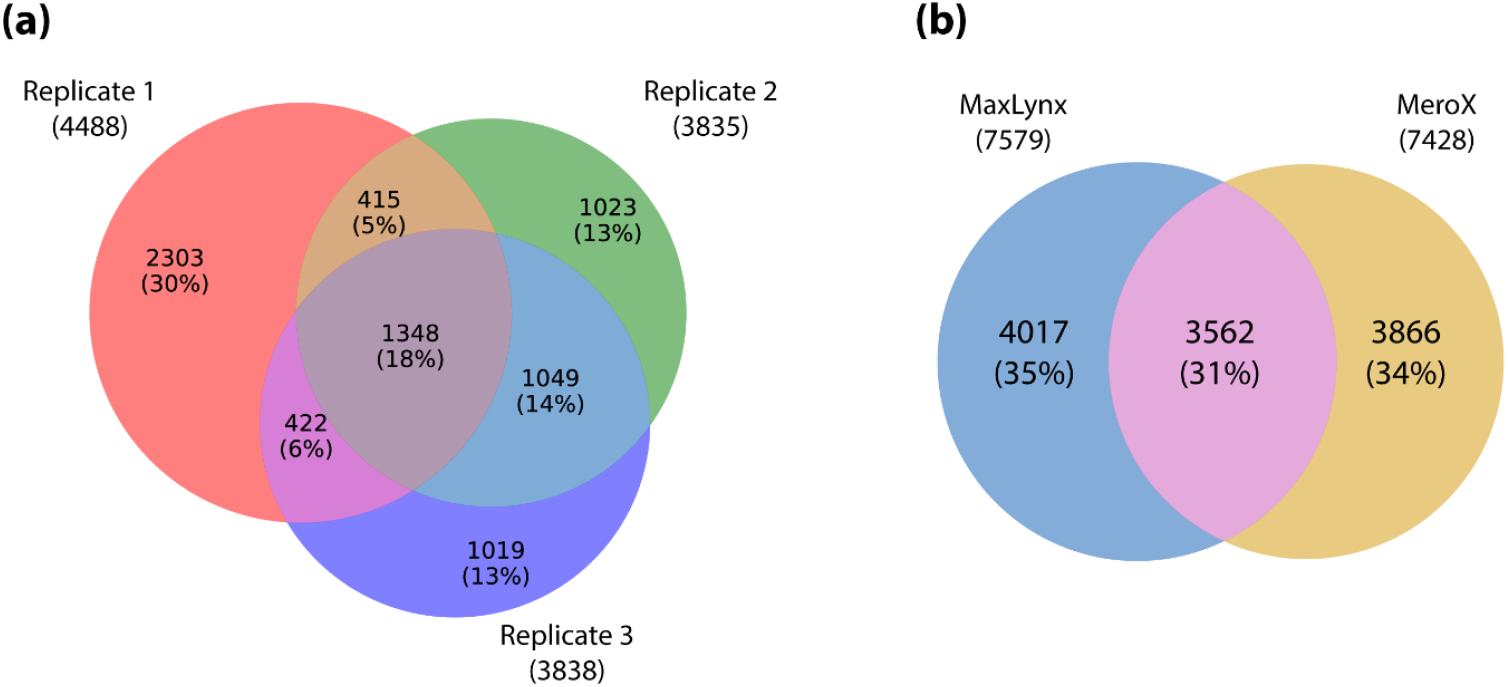
Overlap of unique cross link sites for each replicate at the large scale proteome-wide cross link search. (a) The large scale cross link experiment was performed in three replicates and the absolute number and the percentages are shown. (b) The total number of unique crosslinks for MaxLynx and MeroX are compared.

### Ion-mobility enhanced data

We have used PFU proteins as an entrapment database to evaluate the tims-TOF Pro data set and the number of wrongly assigned CSMs were evaluated, i.e. any CSMs that do not come from intra-BSA protein cross links. The number of CSMs in total were 234 and 226 for DSBU and DSSO data sets, respectively and within these only one and two CSMs were assigned to BSA-PFU protein cross-links, while no PFU-PFU links were found (Table 2). The number of incorrect CSMs are consistent with the CSM-FDR of 1% that was applied to the data. These results show that the search strategy by MaxLynx is sensitive while finding mostly correct hits. Next, we checked how the CCS values behave as a function of molecule mass with respect to the different cross-linked product types (Figure 6). We observe that cross-linked peptides tend to be have higher CCS values along with their high charge states and higher masses compared to linear peptides.

**Table 2.** The overview of the results from the timsTOF Pro the BSA data sets at FDR=1%. The number of CSMs and their unique cross links were reported for both DSBU and DSSO data sets including the CSMs to only BSA intra-protein cross links, in addition to the number of unique cross links.

| Data set | #CSMs (*All*) | # CSMs for (*BSA-BSA*) | #unique cross links (*All*) | #unique cross links (*BSA-BSA*) |
| --- | --- | --- | --- | --- |
| DSBU | 234 | 233 | 126 | 125 |
| DSSO | 226 | 224 | 114 | 112 |

**Figure 6.**
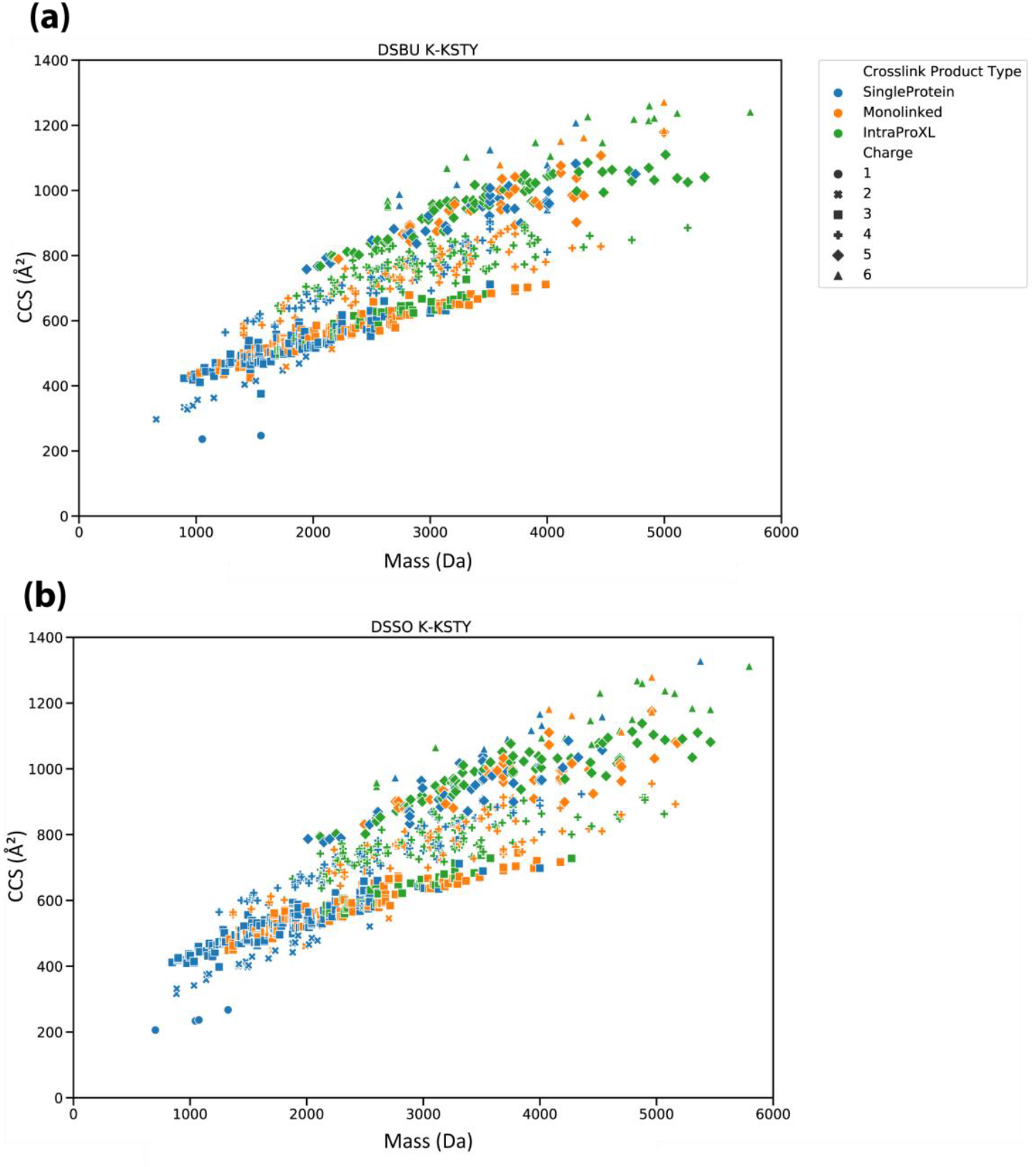
CCS values for TIMS-TOF data set. CCS values are plotted against the molecule mass.

## Conclusion

MaxLynx is a new computational workflow for XL-MS that is integrated into the MaxQuant software. Here we showed that MaxLynx outperforms the benchmarked software for both non-cleavable and MS-cleavable cross-linked peptide data sets at FDR=1%. It works well for data with ion-mobility dimension as well. The success of the MaxLynx was also owing to the peak improvement for heavier peptides like di-peptides. The percentage overlap of cross-links between the replicates is not yet ideal but this may be overcome by better acquisition strategies, and further improvements, such as introducing match-between-runs for cross-linked peptides and applying data-independent acquisition for such samples.

## ASSOCIATED CONTENT

### Supporting information

A step by step guideline how to set up MaxLynx and the extended search settings along with the detailed results for the synthetic cross-linked peptides data sets were provided.

## AUTHOR INFORMATION

### Notes

The authors have declared a potential conflict of interest regarding to this work: F. Busch and N. Nagaraj are employees of Bruker.

## ACKNOWLEDGEMENTS

This project has received funding from the European Union’s Horizon 2020 research and innovation programme under the Marie Skłodowska-Curie grant agreement No 792536 (ŞY). We would like to thank F. Galluzzi, B. Steigenberger and B. Wu for their feedback while testing MaxLynx versions.

## SUPPORTING INFORMATION

### A user guide on how to run MaxLynx

MaxLynx was integrated into the MaxQuant environment. You can download MaxQuant from https://maxquant.org/maxquant/

You must make sure to install .NET Core 2.1 (not higher or not earlier releases) SDK x64 from https://dotnet.microsoft.com/download/dotnet/2.1

#### 1. Load your raw files and set your experiment design

Note that MaxLynx currently works on only one parameter group.

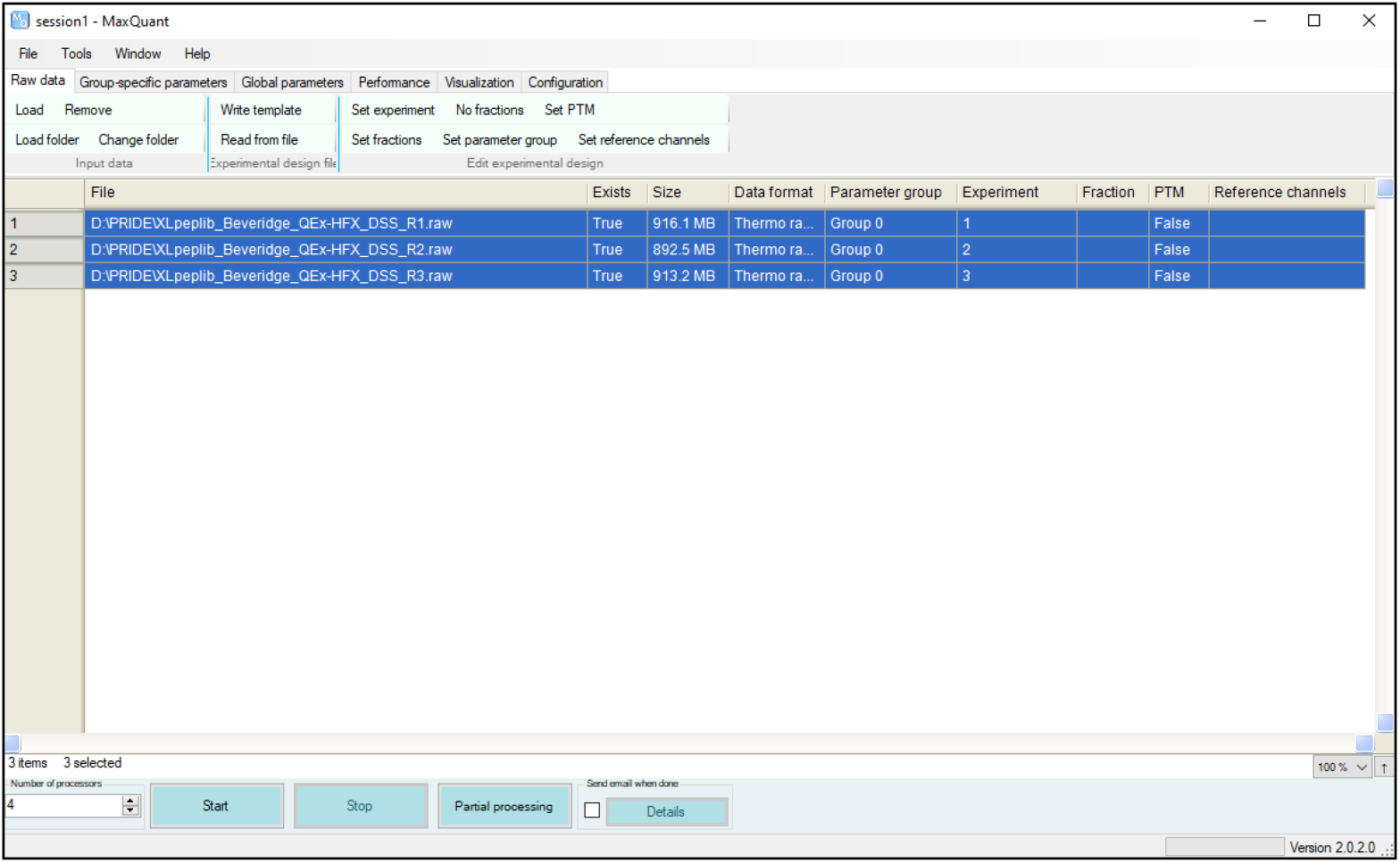

#### 2. Set your MaxLynx parameters on the *Cross links* group-specific parameter tab

This is a new MaxQuant group-specific parameter tab to set MaxLynx runs. It is important to choose the right cross linker type of your experiment.

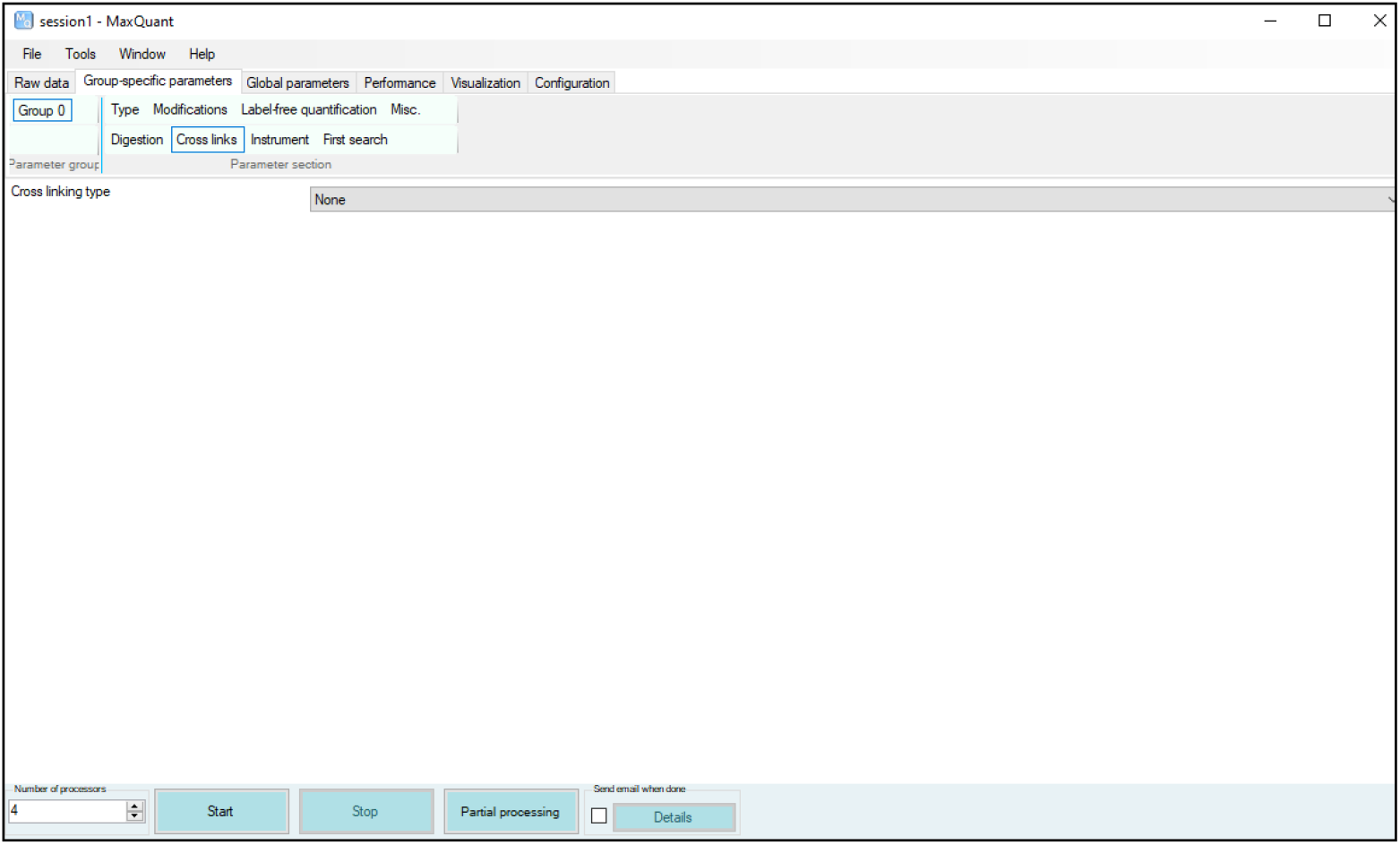

##### 2. a. Enable non-cleavable cross link search

Select by the *Cross linking type* as *Non-cleavable*.

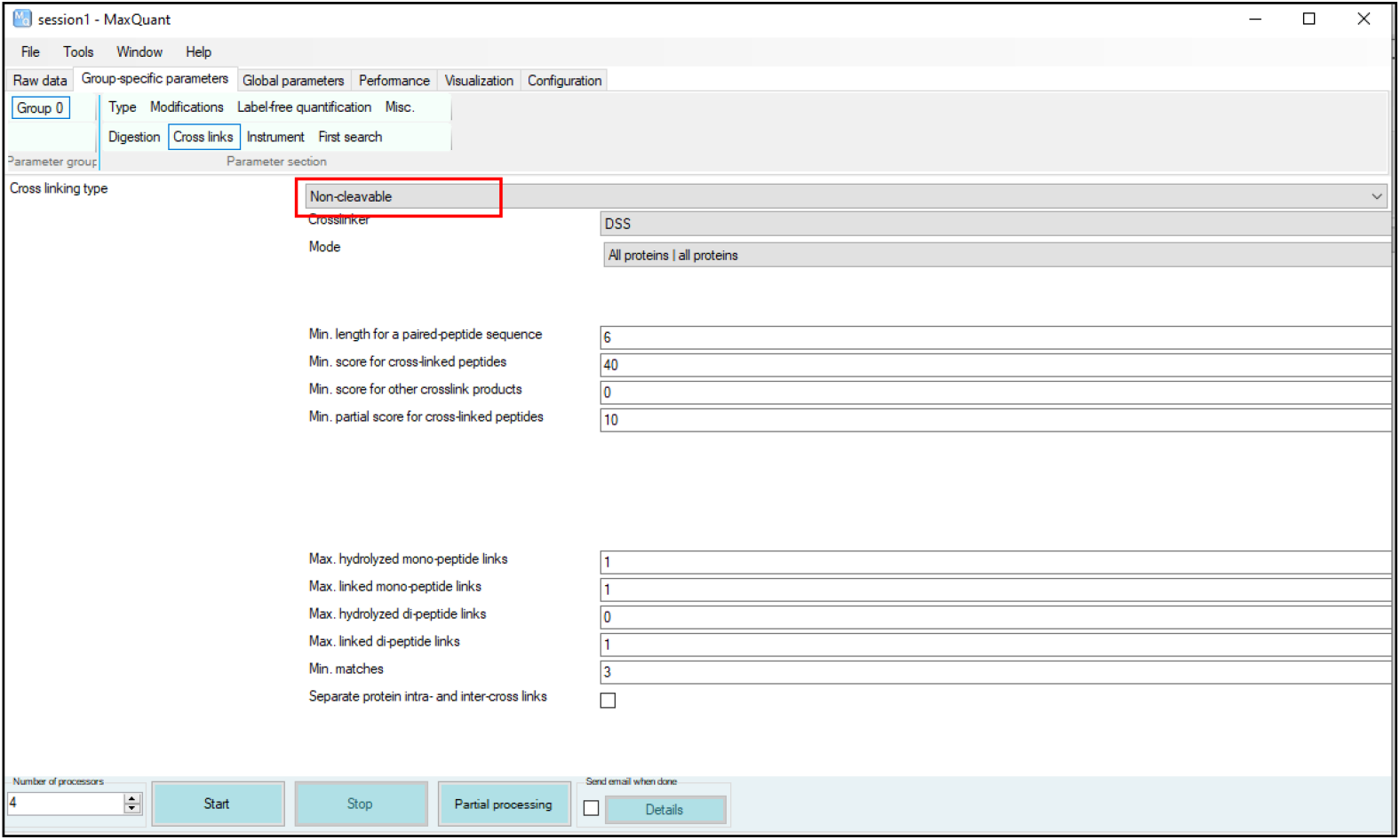

##### 2. b. Enable MS-cleavable cross link search on MS2 level

Select by the *Cross linking type* as *MS2-cleavable*.

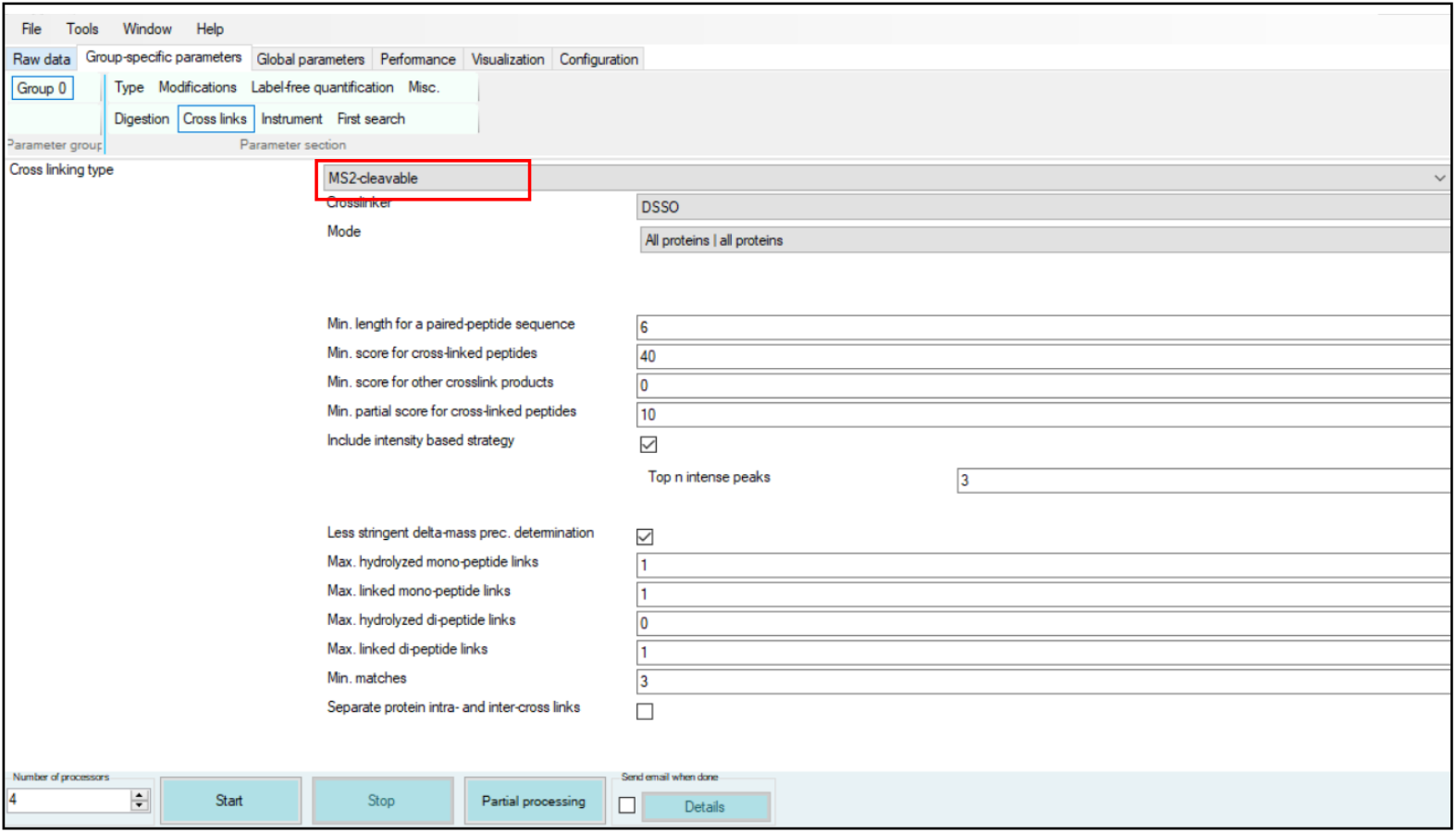

##### 2. c. Configure your cross linkers

You can configure any bi-functional cross linker through this new MaxQuant configuration tab called the *Crosslinks* under the *Configuration* panel. Provide a name and description of your cross linker. Add the compositions for cross link products, where two arms of a cross linker are attached to either mono- or di-peptides (*Linked composition*) and mono-link products, where only one arm of a cross linker is attached to one peptide (*Hydrolyzed composition)* along with the cross linker specificities. You can define *MS-cleavable cross linkers* by selecting the *MS-cleavable* box and then defining the compositions for the long and short parts of this cross linker after cleavage.

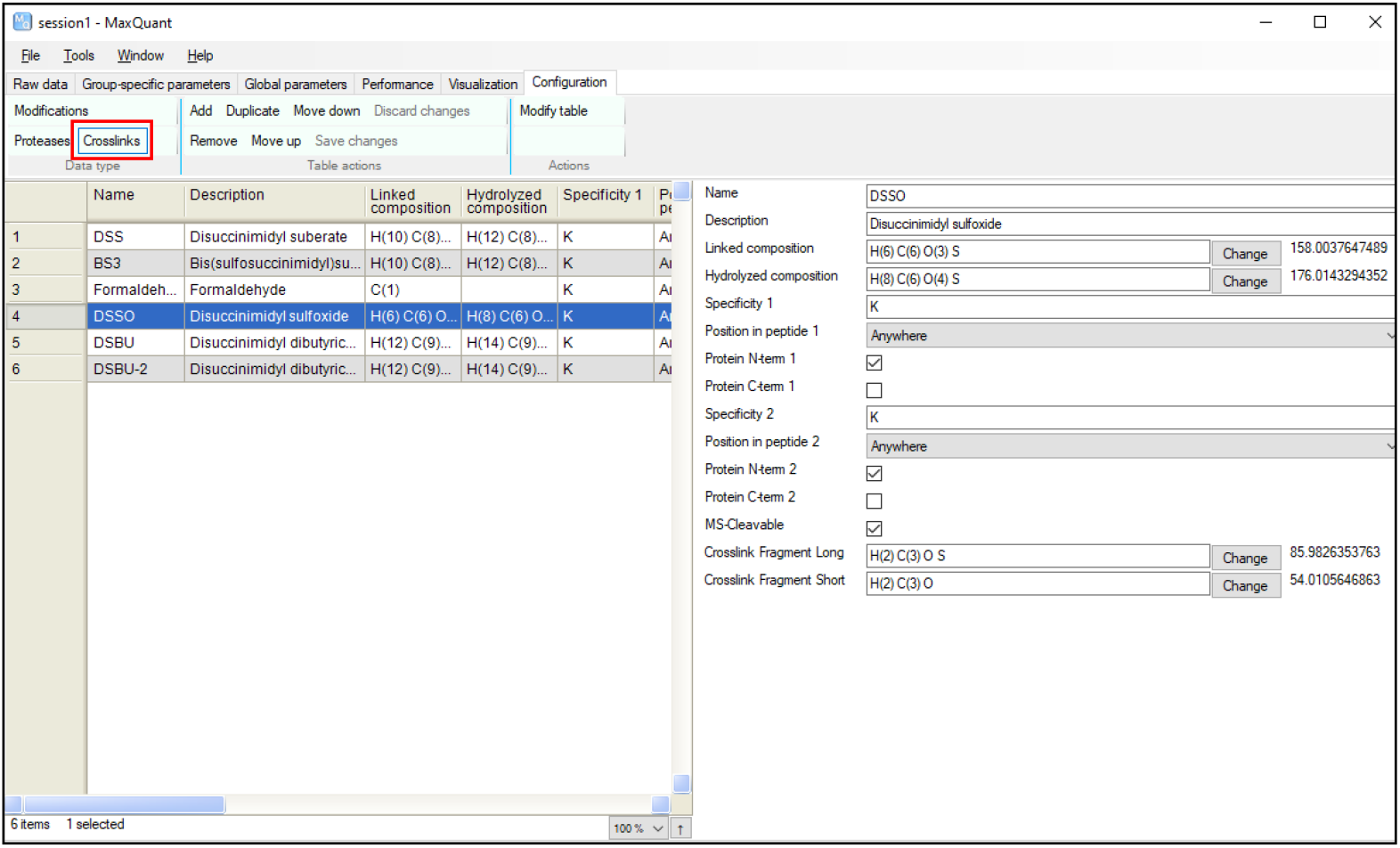

#### 3. Increase the maximum number of missed cleavages and choose the enzyme of your choice

In cross-linked peptide analysis, we tend to change this value to typically three, while the default MaxQuant value is 2 for the maximum missed cleavages

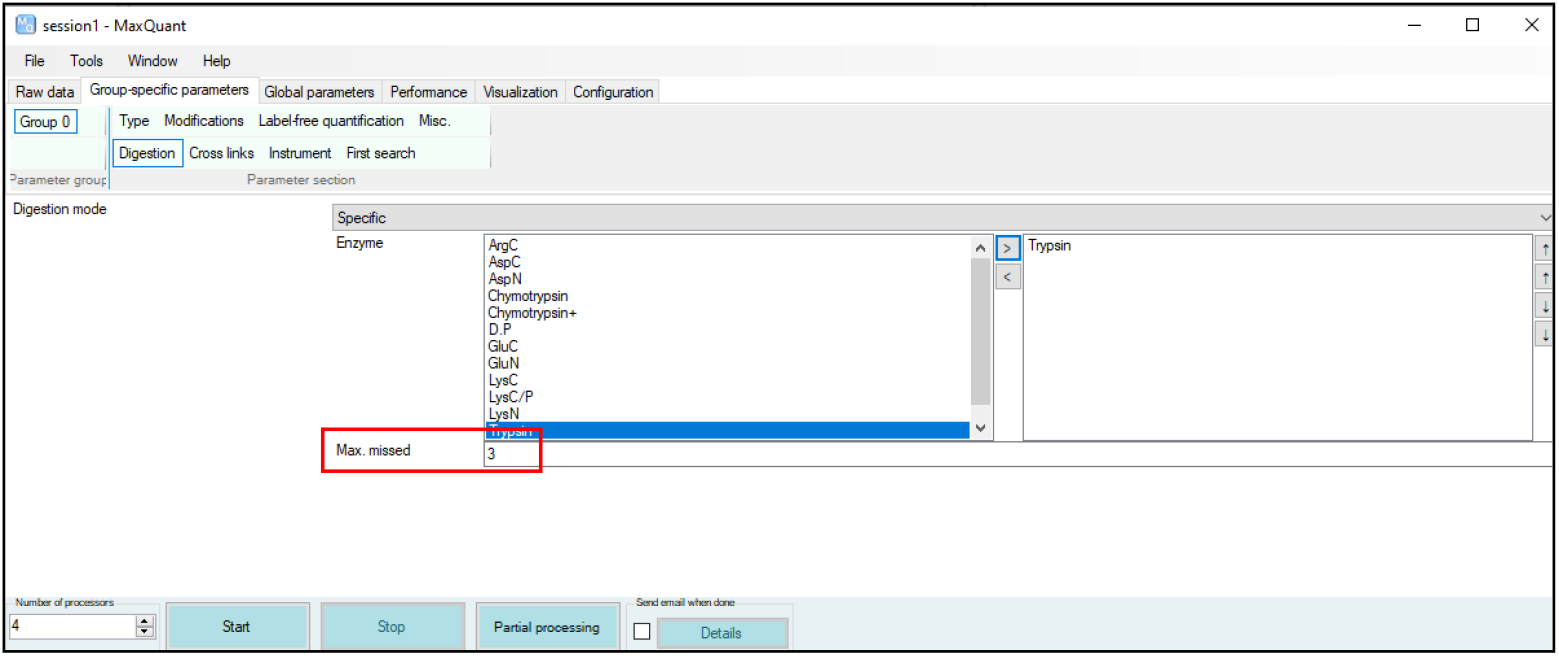

#### 4. Turn on Peak refinement

This new option can be enabled on *Misc*. tab under the *Group-specific parameters* tab.

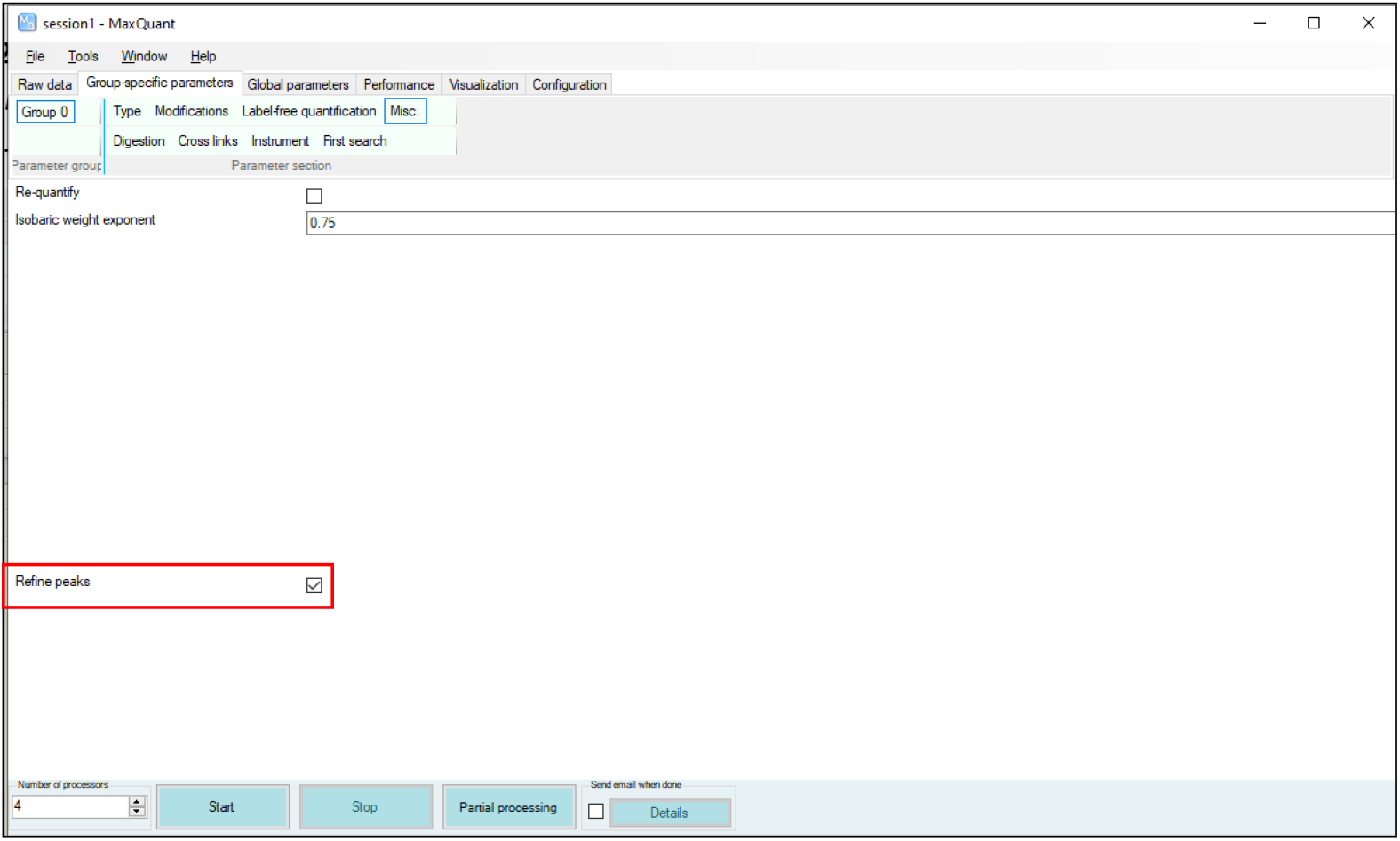

#### 5. Set up protein sequence related information

Add your fasta file but disable including contaminants. Decrease the peptide length to of your choice and increase the peptide mass to adapt cross-linked peptide searches (cross link software typically uses 6000 Da)

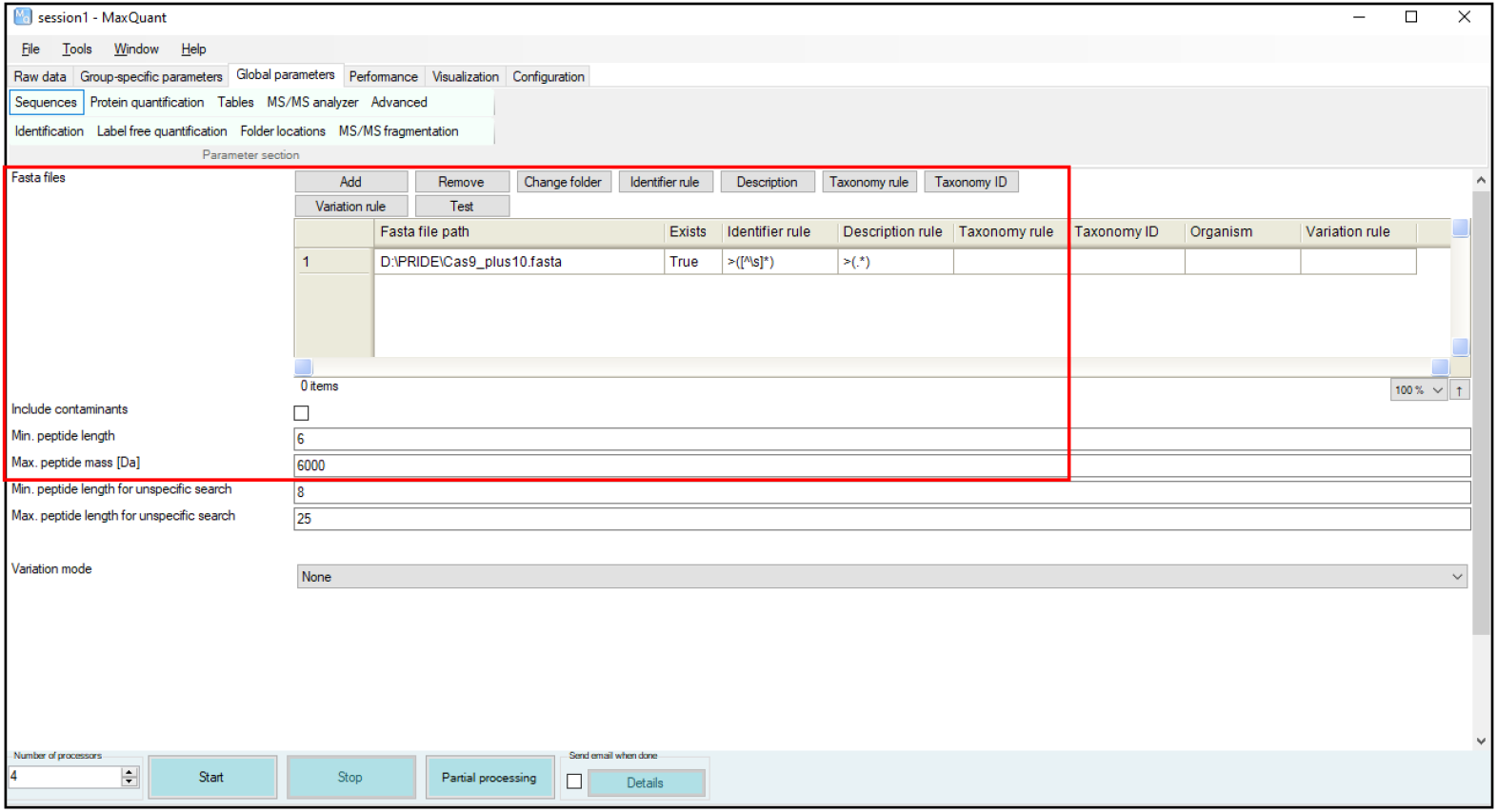

#### 6. Set your FDR values

Here you can currently disable “Second peptide” searches, this option is not functional for this current version.

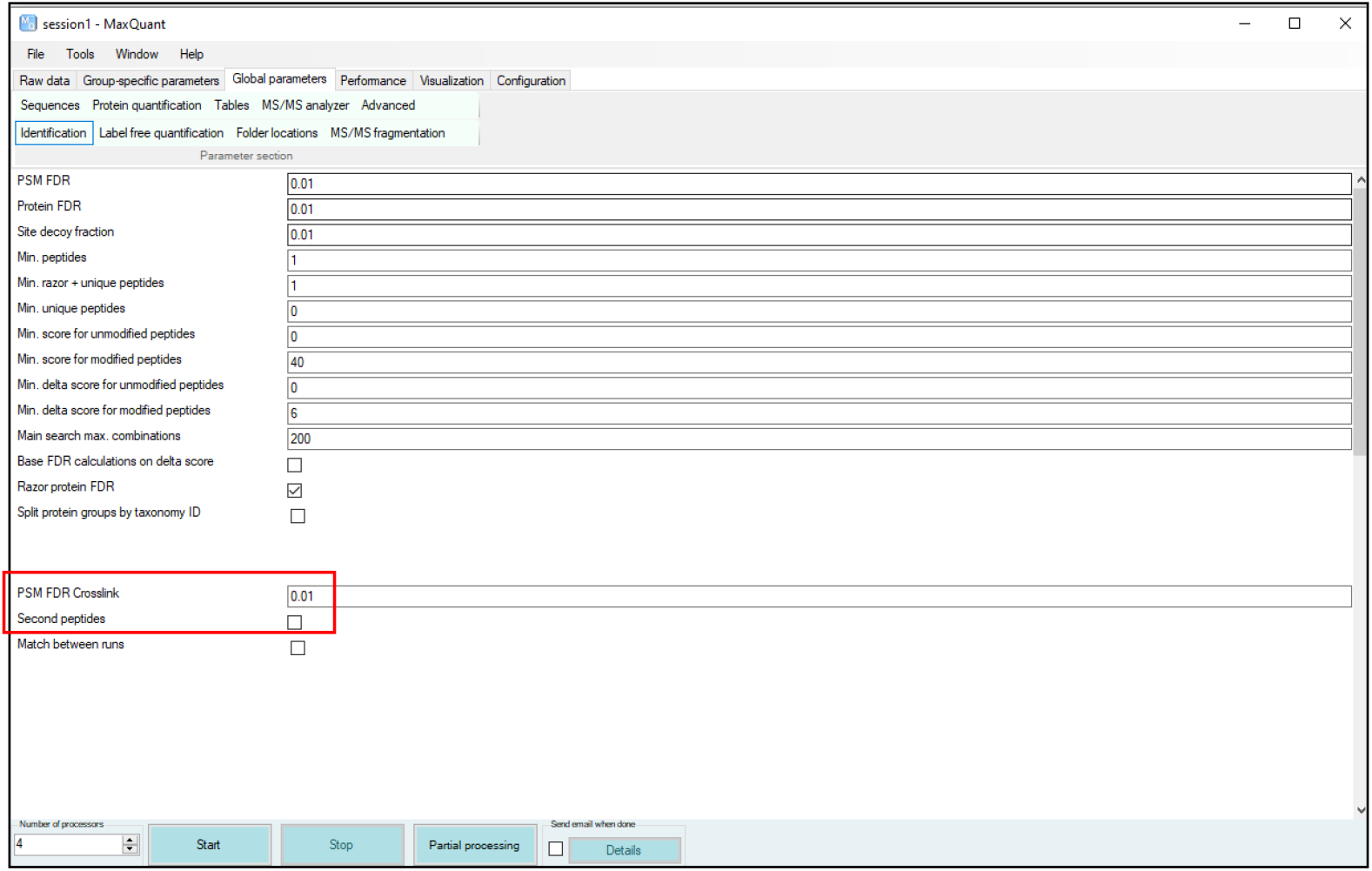

#### 7. Note for Bruker TIMS instruments

Increase the max charge from 4 to 6 because cross-linked peptides tend to have higher charge states.

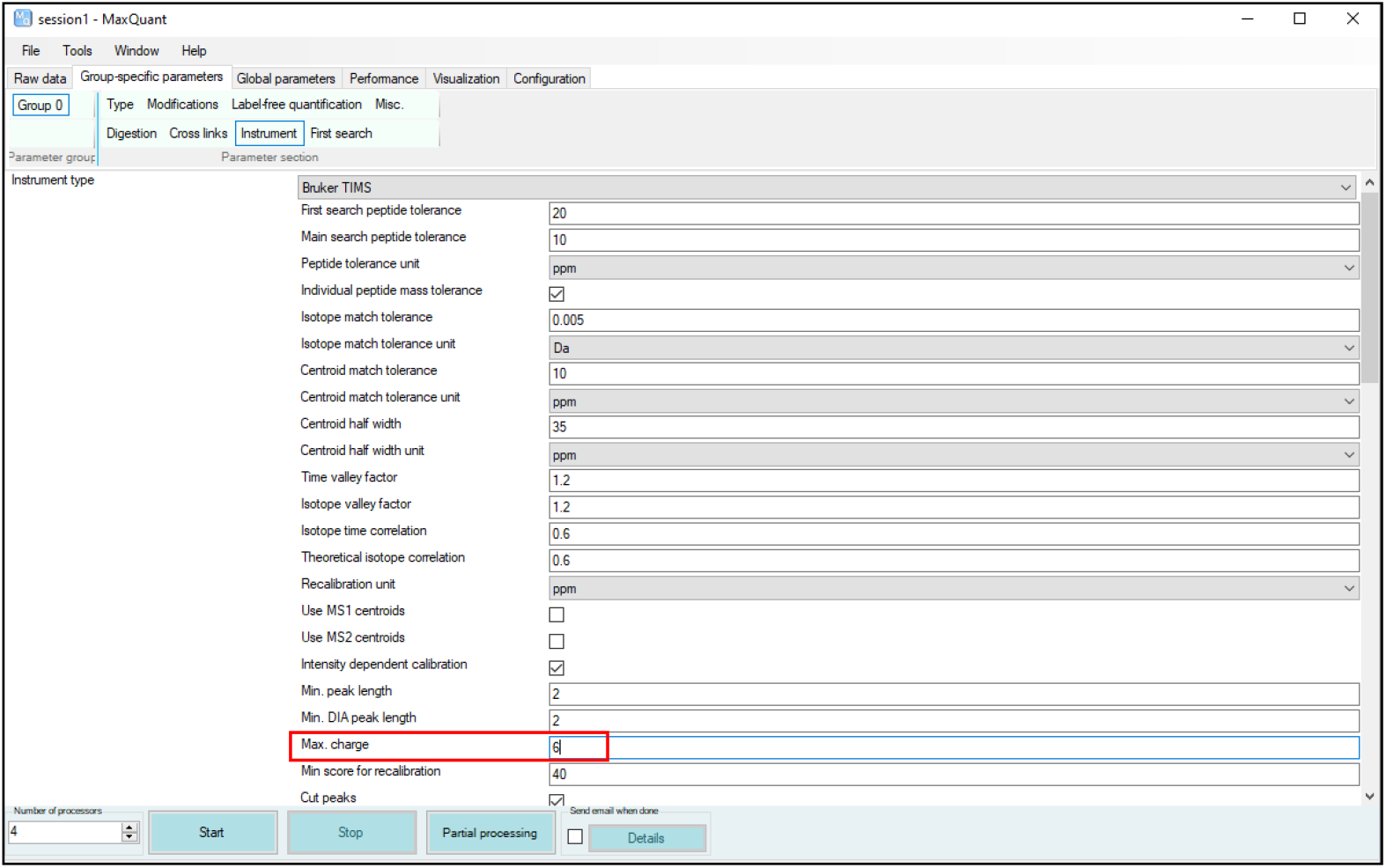

#### 8. Inspect your results

Go to Crosslink MS/MS table (where the information comes from the **crosslinkMsms.txt** table under the **combined/txt** folder after MaxQuant/MaxLynx analysis is finished). Select a row and make sure to be *MS/MS spectra* panel on the visualization. Then click on “*Display selected spectrum”* to view your identification. You can see the peptide sequence-based information on the *Peptide sequence* window under the *MS/MS spectra* panel

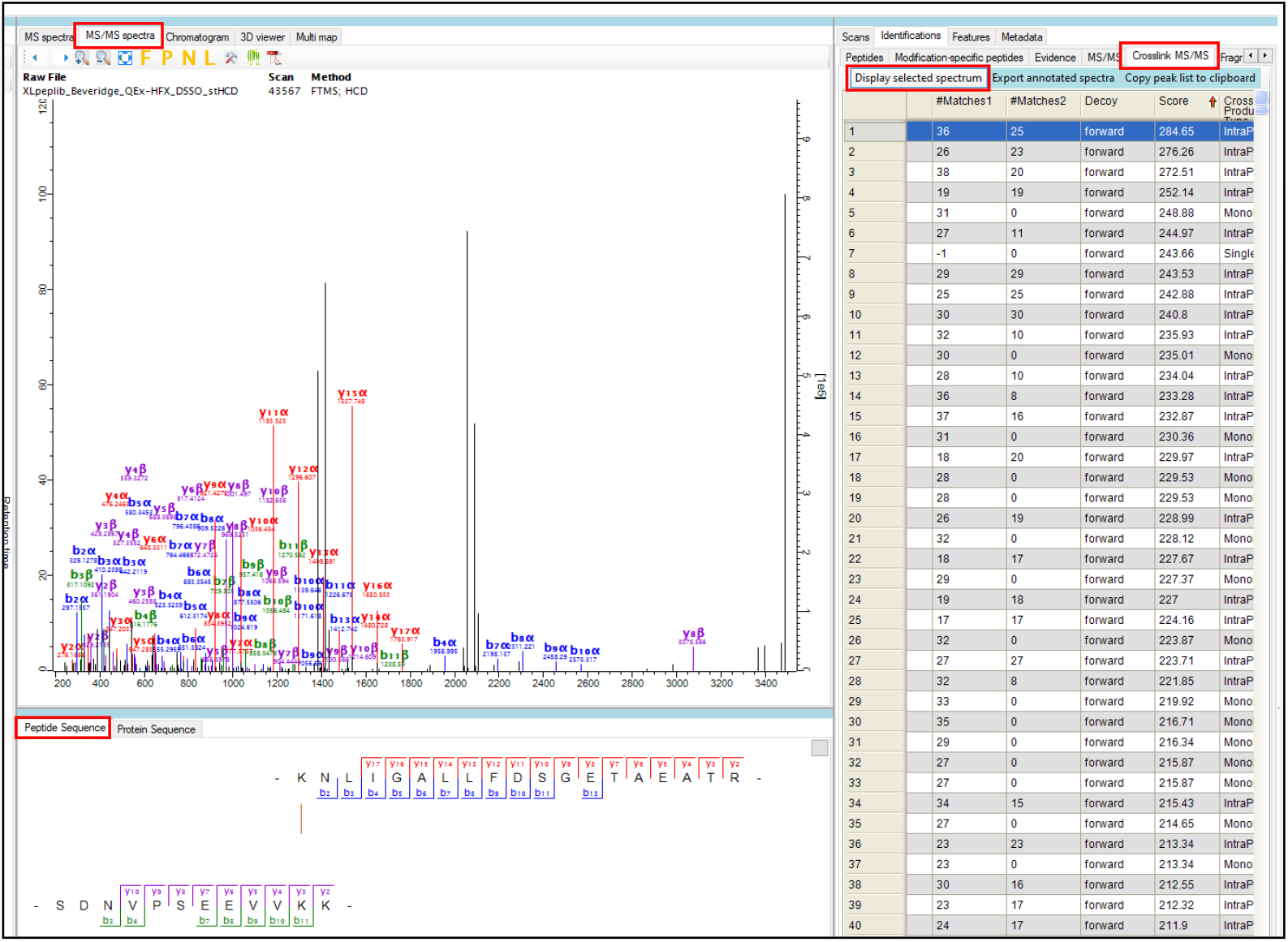

**Supplementary Figure S1.**
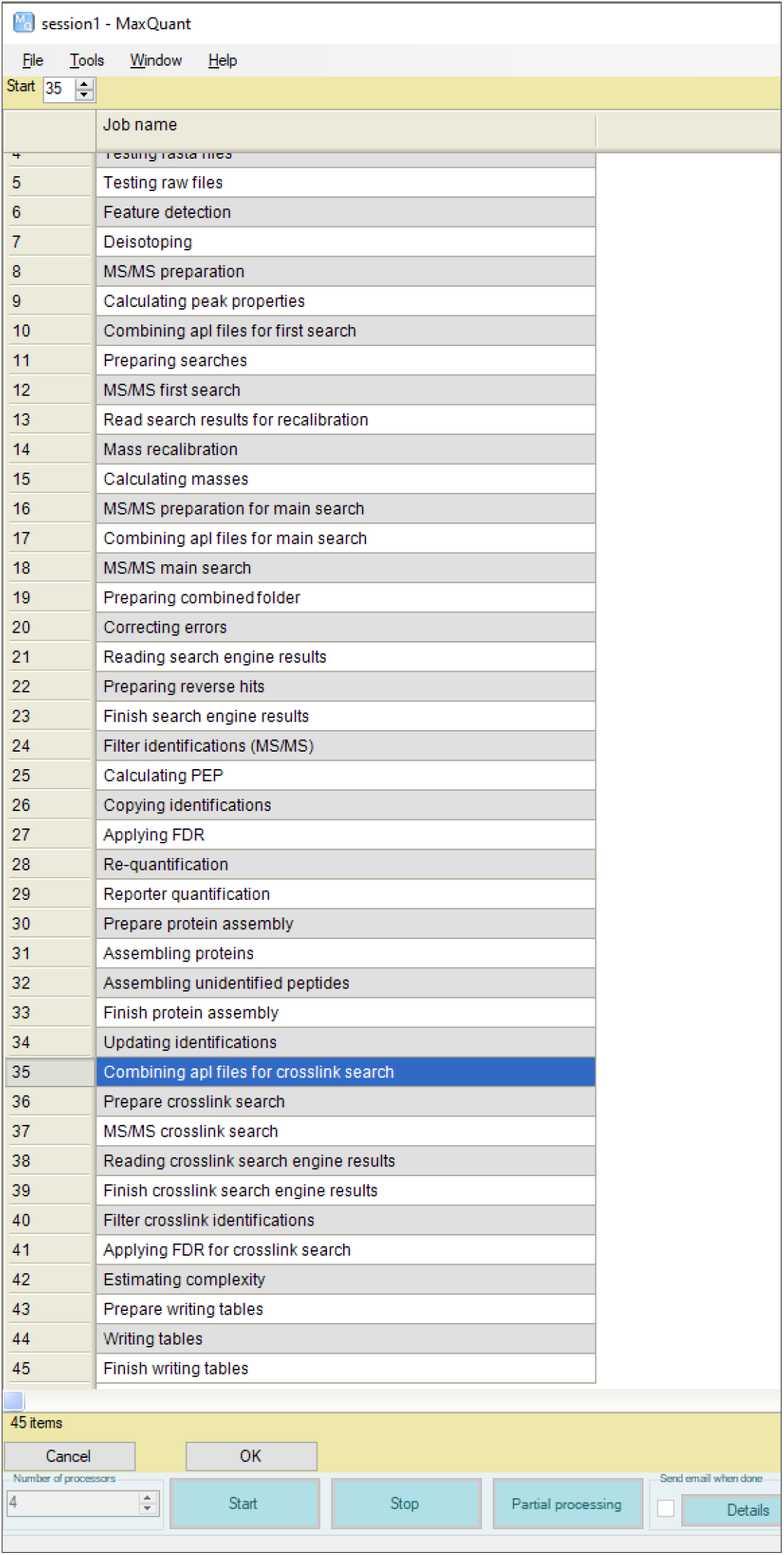
MaxLynx workflow for cross link search

**Supplementary Figure S2.**
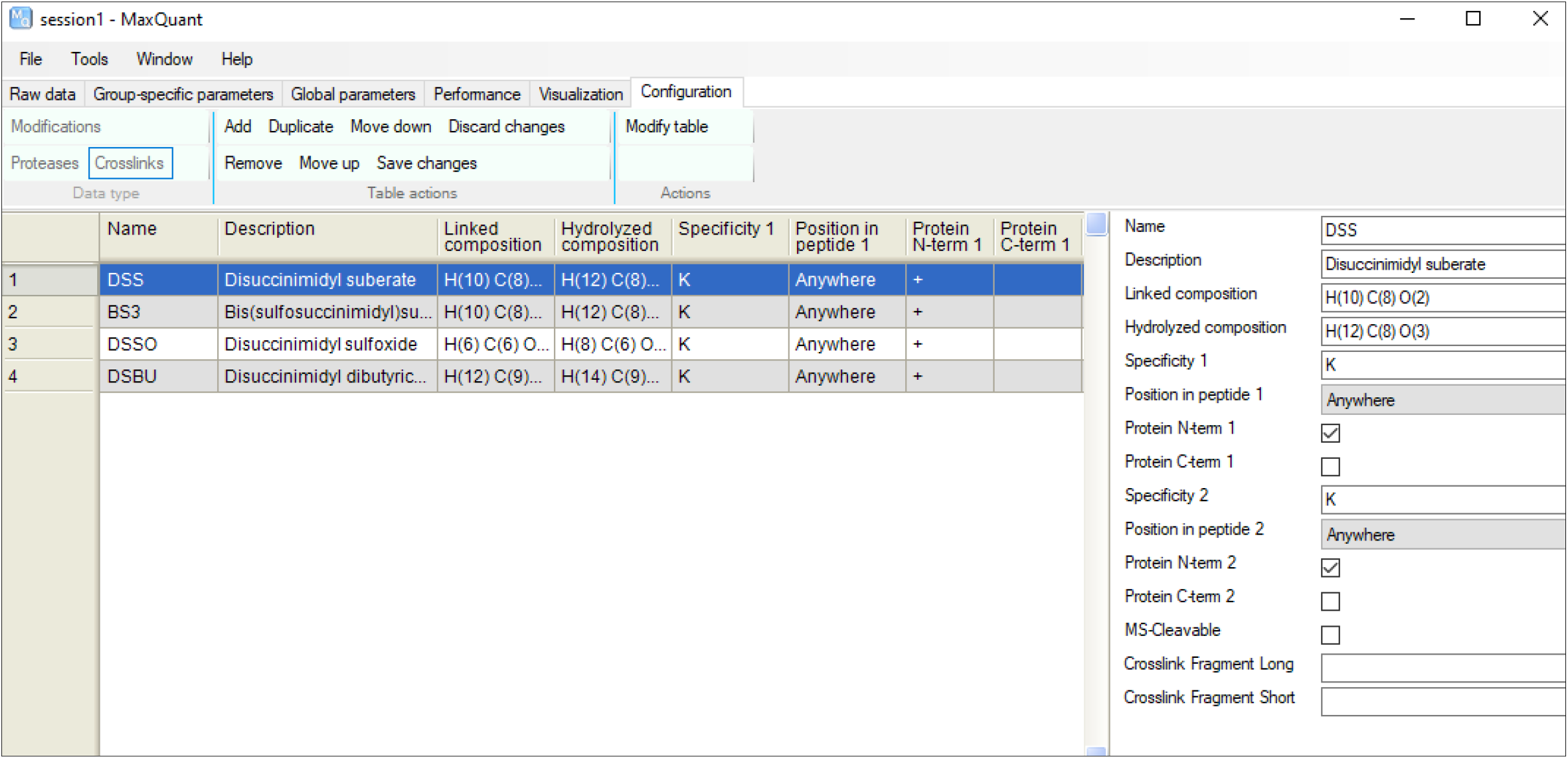
Crosslinks panel. User can easily configure a cross linker of interest through using the Crosslinks panel on the Configuration. See **<u>Step 2.c</u>** on how to run MaxLynx guideline for the further details.

**Supplementary Figure S3.**
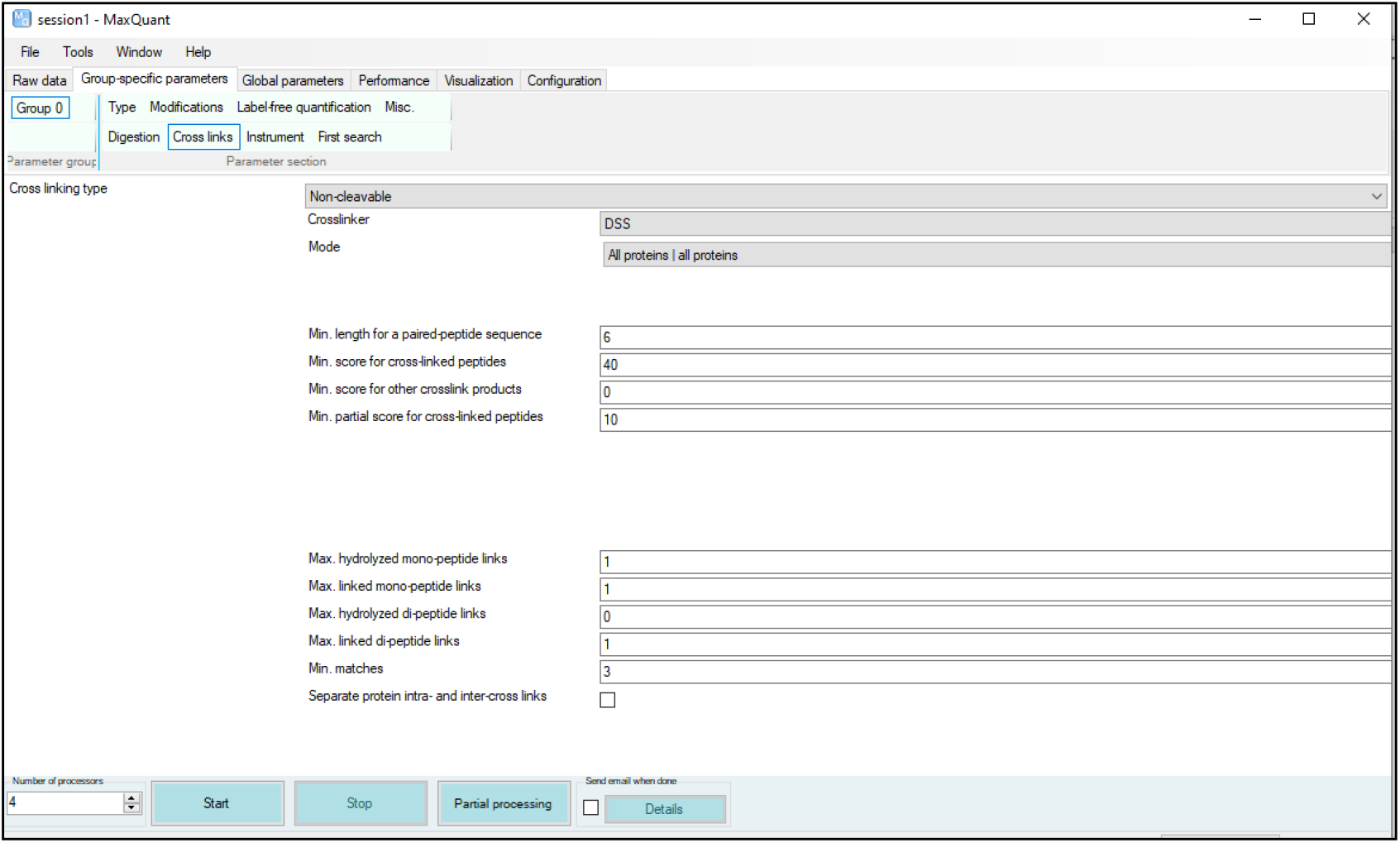
New group-specific parameter panel for cross link searches.

**Supplementary Table S1.** The search settings used for the non-cleavable DSS data set. All the settings except for OpenPepXL and MaxLynx were taken from Beveridge and co-workers^1^, whereas the OpenPepXL settings were from Netz and co-workers^2^. The MaxLynx settings were appended.

|  | **MaxLynx**<br>**(MaxQuant**<br>**v2.0.2)** | **OpenPepX**<br>**L**<br>**(1.1)** | **pLink (2.3.5)** | **StavroX**<br>**(3.6.0)** | **Xi**<br>**(1.6.751)** | **Kojak (1.6.1)** |
| --- | --- | --- | --- | --- | --- | --- |
| **Cross link mass/**<br>**Da** | 138.068 | 138.068 | 138.068 | 138.068 | 138.068 | 138.068 |
| **Mono link**<br>**mass/ Da** | 156.079 | 156.079 | 156.079 | 156.079 | 156.079 | 156.079 |
| **Cross linker**<br>**reactivity** | K-K | K-K | K-K | K-K | K-K | K-K |
| **Fixed**<br>**modification** | Carbamido-<br>methyl [C] | Carbamido-<br>methyl [C] | Carbamido-<br>methyl [C] | Carbamido-<br>methyl [C] | Carbamido-<br>methyl [C] | Carbamido-<br>methyl [C] |
| **Variable**<br>**modification** | Oxidation<br>[M] | Oxidation<br>[M] | Oxidation [M] | Oxidation<br>[M] | Oxidation<br>[M] | Oxidation [M] |
| **Enzyme** | Trypsin | Trypsin | Trypsin | Trypsin | Trypsin | Trypsin |
| **Max. missed**<br>**cleavages** | 3 | 4 | 3 | R:3 K:3 | 3 | 3 |
| **Min peptide**<br>**mass** | - | NA | 500 | 500 | - | 500 |
| **Max peptide**<br>**mass** | 6000 | NA | 6000 | 6000 | - | 6000 |
| **Min peptide**<br>**length** | 5 | 5 | 5 | 5 | 5 | - |
| **Max peptide**<br>**length** | - | NA | 60 | - | - | - |
| **MS1 tolerance**<br>**(ppm)** | 5 | 6 | 5 | 5 | 5 | 5 |
| **MS2 tolerance**<br>**(ppm)** | 20 | 20 | 20 | 20 | 20 | Bin size 0.03<br>Thomson |
| **FDR calculation** | inbuilt | TOPP tool<br>XFDR | inbuilt | inbuilt | Xi FDR<br>(1.1.27) | Percolator<br>(3.02) |
| **FDR level** | PSM | PSM | PSM | PSM | PSM | PSM |

**Supplementary Table S2.** The search settings used for the MS-cleavable data sets. All the settings except for MaxLynx were taken from Beveridge and co-workers^1^ and the MaxLynx settings were appended.

|  |  | **MaxLynx**<br>**(MaxQuant v2.0.2)** | **XlinkX in proteome**<br>**discoverer 2.3** | **MeroX 2.0 beta 5** |
| --- | --- | --- | --- | --- |
| **DSBU** | **Cross link**<br>**mass/ Da** | 196.085 | 196.085 | 196.085 |
|  | **Bu-**<br>**fragment/ Da** | 85.053 | - | 85.053 |
|  | **BuUr-**<br>**fragment/ Da** | 111.032 | - | 111.032 |
| **DSSO** | **Cross link**<br>**mass/ Da** | 158.004 | 158.004 | 158.004 |
|  | **Alkene/ Da** | 54.011 | - | 54.011 (essential) |
|  | **Thiol/ Da** | 85.983 | - | 85.983 (essential) |
|  | **Sulfenic acid/**<br>**Da** | - | - | 103.993 |
| **Cross linker reactivity** |  | K-K | K-K | K-K |
| **Fixed modification** |  | Carbamidomethyl [C] | Carbamidomethyl [C] | Carbamidomethyl [C] |
| **Variable modification** |  | Oxidation [M] | Oxidation [M] | Oxidation [M] |
| **Enzyme** |  | Trypsin | Trypsin | Trypsin |
| **Max. missed**<br>**cleavages** |  | 3 | 3 | R:3 K:3 |
| **Min peptide mass** |  | - | 500 | 500 |
| **Max peptide mass** |  | 6000 | 6000 | 6000 |
| **Min peptide length** |  | 5 | 5 | 5 |
| **MS1 tolerance (ppm)** |  | 5 | 5 | 5 |
| **MS2 tolerance (ppm)** |  | 20 | 20 | 20 |
| **S/N ratio** |  | - | 1.5 | 1.5 |
| **FDR calculation** |  | inbuilt | inbuilt | inbuilt |
| **FDR level** |  | PSM | PSM | PSM |

**Supplementary Table S3.**
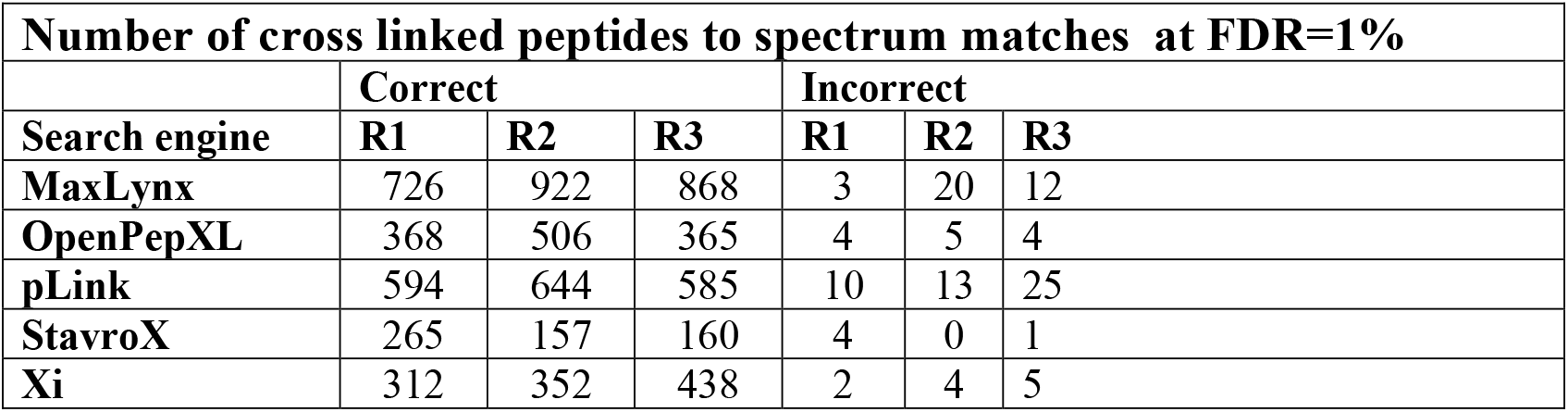
Number of cross linked peptides to spectrum matches (CSMs) in the DSS data set at **1% FDR**. The OpenPepXL results were taken from Netz and co-workers ^2^ whereas the results of pLink, StavroX and Xi from Beveridge and co-workers ^1^.

**Supplementary Table S4.**
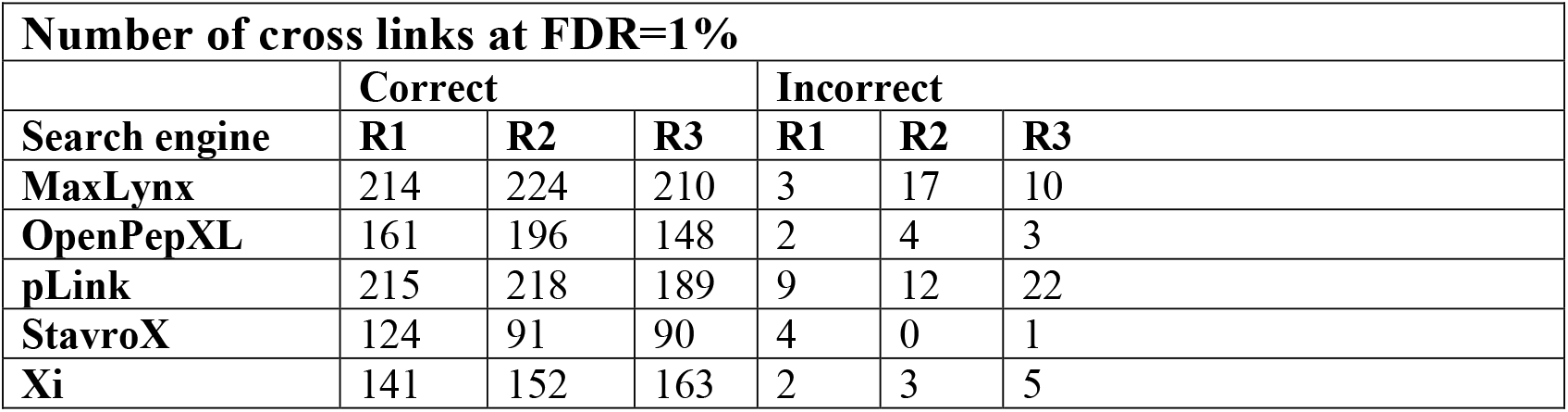
Number of unique cross links in the DSS data set at **1% FDR**. The OpenPepXL results were taken from Netz and co-workers ^2^ whereas the results of pLink, StavroX and Xi from Beveridge and co-workers ^1^.

**Supplementary Table S5.** The number of cross links identified on the MS-cleavable cross linker data sets, DSBU and DSSO respectively at **1% FDR**. MeroX and XlinkX results were taken from the Beveridge and co-workers^1^.

| Number of cross links at FDR 1% |  |  |
| --- | --- | --- |
| Cross linker, Search engines | Correct | Incorrect |
| DSBU, MaxLynx | 240 | 12 |
| DSBU, MeroX, Rise | 207 | 11 |
| DSBU, MeroX, Riseup | 237 | 15 |
| DSBU, XlinkX | 120 | 37 |
| DSSO, MaxLynx | 185 | 10 |
| DSSO, MeroX, Rise | 124 | 1 |
| DSSO, MeroX, Riseup | 149 | 19 |
| DSSO, XlinkX | 128 | 53 |

